# Environmental and genetic contributions to imperfect *w*Mel-like *Wolbachia* transmission and frequency variation

**DOI:** 10.1101/2020.06.09.142828

**Authors:** Michael T.J. Hague, Heidi Mavengere, Daniel R. Matute, Brandon S. Cooper

## Abstract

Maternally transmitted *Wolbachia* bacteria infect about half of all insect species. They usually show imperfect maternal transmission and often produce cytoplasmic incompatibility (CI). Irrespective of CI, *Wolbachia* frequencies tend to increase when rare only if they benefit host fitness. Several *Wolbachia*, including *w*Mel that infects *Drosophila melanogaster* cause weak or no CI and persist at intermediate frequencies. On the island of São Tomé off West Africa, the frequencies of *w*Mel-like *Wolbachia* infecting *D. yakuba* (*w*Yak) and *D. santomea* (*w*San) fluctuate, and the contributions of imperfect maternal transmission, fitness effects, and CI to these fluctuations are unknown. We demonstrate spatial variation in *w*Yak frequency and transmission on São Tomé. Concurrent field estimates of imperfect maternal transmission do not predict spatial variation in *w*Yak frequencies, which are highest at high altitudes where maternal transmission is the most imperfect. Genomic and genetic analyses provide little support for *D. yakuba* effects on *w*Yak transmission. Instead, rearing at cool temperatures reduces *w*Yak titer and increases imperfect transmission to levels observed on São Tomé. Using mathematical models of *Wolbachia* frequency dynamics and equilibria, we infer temporally variable imperfect transmission or spatially variable effects on host fitness and reproduction are required to explain *w*Yak frequencies. In contrast, spatially stable *w*San frequencies are plausibly explained by imperfect transmission, modest fitness effects, and weak CI. Our results provide insight into causes of *w*Mel-like frequency variation in divergent hosts. Understanding this variation is crucial to explain *Wolbachia* spread and to improve *w*Mel biocontrol of human disease in transinfected mosquito systems.

## INTRODUCTION

Maternally transmitted *Wolbachia* bacteria are the most widespread group of intracellular symbionts, infecting about half of all insect species, as well as other arthropods and nematodes (Werren *et al.* 2008; Zug and Hammerstein 2012; Weinert *et al.* 2015). *Wolbachia* often manipulate host reproduction to favor infected females (Rousset *et al.* 1992; Hoffmann and Turelli 1997; Hurst and Jiggins 2000). For example, *Wolbachia* like *w*Ri that infects *Drosophila simulans* cause strong cytoplasmic incompatibility (CI) that reduces egg hatch when uninfected females mate with *Wolbachia*-infected males (Hoffmann and Turelli 1997)—infected females are protected from CI (Shropshire *et al.* 2018), providing them with a relative fitness advantage. Many other *Wolbachia* do not strongly manipulate host reproduction yet persist in nature. These include *w*Mau in *D. mauritiana* (Meany *et al.* 2019), *w*Au in *D. simulans* (Hoffmann *et al.* 1996), *w*Suz in *D. suzukii* (Hamm et al. 2014; Cattel et al. 2016), *w*Mel in *D. melanogaster* (Kriesner *et al.* 2016), and *Wolbachia* variants infecting *D. yakuba*-clade hosts (*w*Yak in *D. yakuba*, *w*San in *D. santomea*, and *w*Tei in *D. teissieri*) that diverged from *w*Mel in only the last 30,000 years (Charlat *et al.* 2004; Zabalou *et al.* 2004; Cooper *et al.* 2017, 2019).

Population frequency dynamics and equilibria of *Wolbachia* can be approximated with three parameters: (1) the proportion of uninfected ova produced by infected females (*μ*; i.e., imperfect maternal transmission), (2) the fitness of infected females relative to uninfected females (*F*; i.e., components of host fitness like fecundity), and (3) the relative egg hatch of uninfected eggs fertilized by infected males (*H*; i.e., the severity of CI) (Hoffmann *et al.* 1990). When initially rare, *Wolbachia* must generate *F*(1-*μ*) > 1 to spread deterministically from low frequencies, regardless of whether they cause CI. The specific fitness benefits underlying low-frequency spread are poorly understood, but potential candidates include fecundity effects (Weeks *et al.* 2007), viral protection (Hedges *et al.* 2008; Teixeira *et al.* 2008; Osborne *et al.* 2009; Martinez *et al.* 2014), and nutrient provisioning (Brownlie *et al.* 2009; Hosokawa *et al.* 2010; Nikoh *et al.* 2014; Moriyama *et al.* 2015). Once infections become sufficiently common, strong CI drives *Wolbachia* to high equilibrium frequencies balanced by imperfect maternal transmission (Hoffmann *et al.* 1990; Barton and Turelli 2011), as observed for rapid *w*Ri spread through global *D. simulans* populations (Turelli and Hoffmann 1991, 1995; Carrington *et al.* 2011; Kriesner *et al.* 2013). Conversely, *Wolbachia* that do not cause strong CI tend to occur at intermediate frequencies that fluctuate through time and space (*w*Suz, Hamm *et al.* 2014; *w*Mel, Kriesner *et al.* 2016; *D. yakuba*-clade *Wolbachia*, Cooper *et al.* 2017), suggesting the parameters influencing *Wolbachia* spread must vary.

Imperfect maternal transmission has been documented for *w*Ri and *w*Au in *D. simulans* and *w*Mel in *D. melanogaster* in the field (Hoffmann *et al.* 1990, 1998; Turelli and Hoffmann 1995; Carrington *et al.* 2011). In contrast, maternal transmission is nearly perfect under laboratory conditions (Hoffmann *et al.* 1990; Meany *et al.* 2019), indicating environmental conditions influence the fidelity of *Wolbachia* transmission. Transmission is predicted to depend on *Wolbachia* titer and localization within developing female oocytes (Ferree *et al.* 2005; Serbus and Sullivan 2007; Casper-Lindley *et al.* 2011; Serbus *et al.* 2015). Host diet influences *Wolbachia* titer and localization during late oogenesis (Serbus *et al.* 2015; Camacho *et al.* 2017; Christensen *et al.* 2019), suggesting that seasonal or spatial availability of preferred fruits (e.g., marula fruit for *D. melanogaster* or figs for *D. santomea*) could potentially affect maternal transmission (Cariou *et al.* 2001; Mansourian *et al.* 2018; Sprengelmeyer *et al.* 2020). Thermal conditions may directly alter *Wolbachia* transmission. For instance, stressful temperatures disrupt *w*Mel transmission in transfected *Aedes aegypti* mosquitoes (Ross *et al.* 2017; Foo *et al.* 2019; Ross *et al.* 2019a), potentially reducing the efficacy of *w*Mel biocontrol of human disease transmission (Hoffmann *et al.* 2011; Van den Hurk *et al.* 2012; Aliota *et al.* 2016; Caragata *et al.* 2016; O’Neill 2018). The frequency of *w*Mel in its natural *D. melanogaster* host declines at temperate latitudes in Eastern Australia and Eastern North America, which may be due to a combination of *Wolbachia* fitness costs and reduced maternal transmission in cold environments (Kriesner *et al.* 2016). Together, these data suggest nutritional and thermal variation may perturb *Wolbachia* maternal transmission, particularly for *w*Mel-like *Wolbachia*.

*w*Mel-like *Wolbachia* infect the sister-species *D. yakuba* and *D. santomea* that diverged from model *w*Mel-infected *D. melanogaster* up to 13 million years ago (Tamura *et al.* 2004). *D. yakuba* is a human commensal distributed throughout sub-Saharan Africa that is generally found in open habitats, but absent in rainforests (Llopart *et al.* 2005a; b; Cooper *et al.* 2018). On Pico de São Tomé off West Africa, *D. yakuba* is found in open disturbed areas below 1,450 m, whereas endemic *D. santomea* is found between 1,153 and 1,800 m in montane mist and rain forests. The distributions of *D. yakuba* and *D. santomea* overlap in the midlands of Pico de São Tomé where they hybridize (Lachaise *et al.* 2000; Comeault *et al.* 2016; Turissini and Matute 2017; Cooper *et al.* 2017). Phylogenomic analyses indicate that *w*Mel-like *Wolbachia* spread among *D. yakuba*-clade host species via introgressive transfer in the last 2,500 to 4,500 years (Cooper *et al.* 2019). *w*Yak and *w*San share very high sequence similarity across their genomes (0.0017% third-position pairwise differences; 643 genes across 644,586 bp; Cooper *et al.* 2019); and like *w*Mel, they cause weak CI potentially modulated by host factors (Reynolds and Hoffmann 2002; Cooper *et al.* 2017). Relative to *w*Mel (Beckmann *et al.* 2017; LePage *et al.* 2017), these *Wolbachia* have an additional set of loci implicated in CI that they acquired horizontally from divergent B-group *Wolbachia* (Cooper *et al.* 2019). Infection frequencies on São Tomé are temporally variable in idiosyncratic ways, such that *w*Yak frequency increased from 2001 (0.40) to 2009 (0.76; *P* < 0.001) and *w*San decreased from 2009 (0.77) to 2015 (0.37; *P* < 0.0001), during a period where *w*Yak frequencies were stable. *w*Yak frequencies in 2009 also varied spatially between São Tomé and the nearby island of Bioko (0.03; *P* < 0.001) (Cooper *et al.* 2017). This variation is similar to sporadic fluctuations of *w*Mel observed in Africa (Kriesner *et al.* 2016), and suggests that *w*Mel-like *Wolbachia* with high sequence similarity may behave differently across different abiotic conditions and host backgrounds.

The causes of *w*Mel-like *Wolbachia* frequency fluctuations remain mostly unknown. Here, we use a combination of field, laboratory, and mathematical analyses to dissect the contributions of imperfect maternal transmission, *Wolbachia* effects on host fitness, and CI to spatiotemporal *w*Yak and *w*San frequency variation on São Tomé. In 2018, we generated concurrent estimates of imperfect maternal transmission (*μ*) and infection frequencies (*p*) of *w*Yak and *w*San along an altitudinal transect on Pico de São Tomé to test whether field estimates of *μ* predict spatial variation in *p*. In the laboratory, we then tested how environmental conditions and host genetic factors contribute to variation in imperfect maternal transmission. Finally, we used mathematical models of *Wolbachia* frequency dynamics and equilibria to better understand *w*Mel-like frequency variation on São Tomé. Our results provide insight into the basis of *w*Mel-like *Wolbachia* frequency fluctuations in two sister species that diverged from model *D. melanogaster* up to 13 million years ago (Tamura *et al.* 2004).

## MATERIALS AND METHODS

### Estimating *Wolbachia* frequencies and imperfect maternal transmission

To determine spatial variation in *w*Yak and *w*San frequencies, we sampled female *D. yakuba* (*N* = 81) and *D. santomea* (*N* = 78) from 15 trapping sites on São Tomé in 2018 (Figure 1; Table S1). Flies were sampled using fruit traps and by sweeping nets over piles of local jackfruit and bananas. Flies sampled from the four *D. yakuba* traps were assigned to a low altitude (*N* = 40) and high altitude group (*N* = 41). The low altitude region (590 m) is characterized by relatively hot and dry conditions, whereas high altitude (900–1,104 m) is generally cooler and more humid. The high altitude sites are contiguous with the known *D. yakuba*-*D. santomea* hybrid zone (Comeault *et al.* 2016; Turissini and Matute 2017). *D. santomea* individuals sampled from eleven traps were initially assigned into two regional groups based on geography such that site 1 (*N* = 24) and site 2 (*N* = 54) were collected on opposite sides of a small mountainous region (Figure 1). Subsequent analyses found no regional variation in *p* or *μ* for *w*San (see below).

**FIGURE 1.**
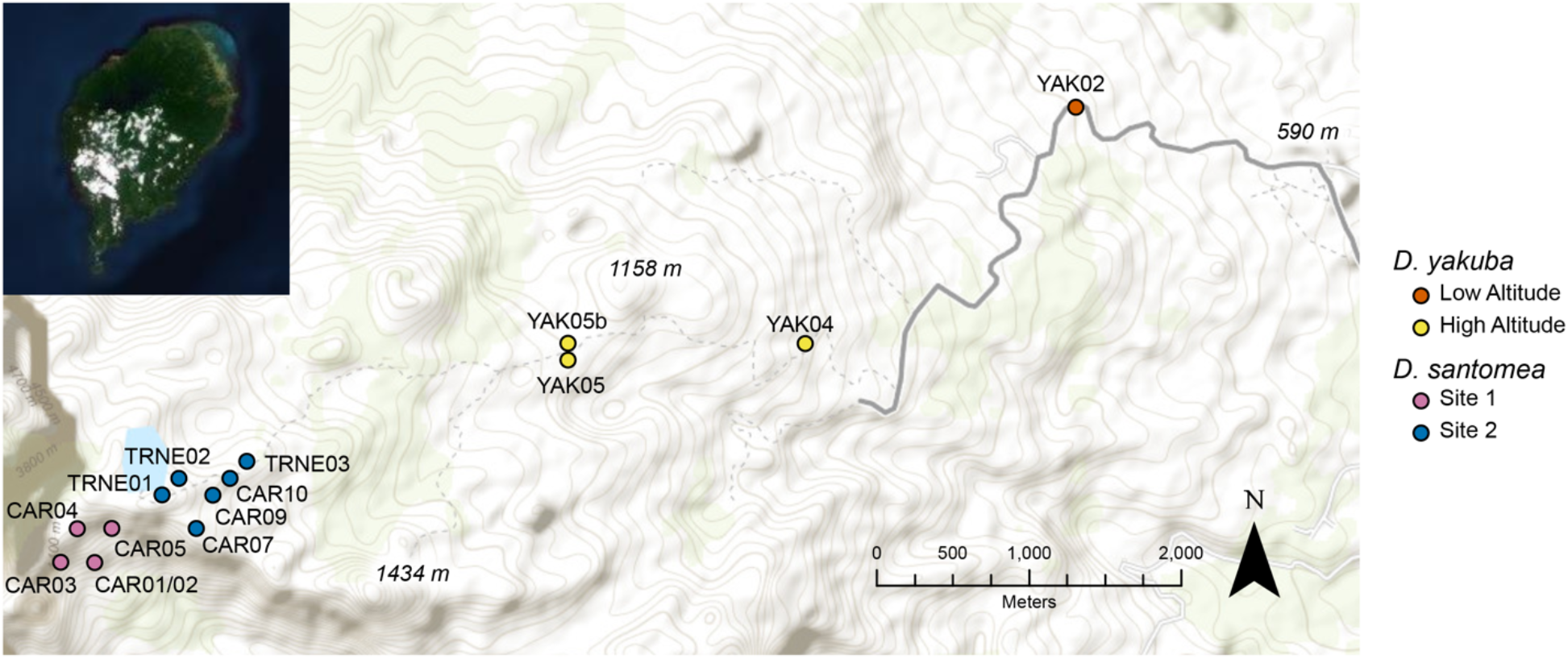
Trapping sites on the island of São Tomé (inset) off West Africa. Points denote individual trapping sites (Table S1), which are grouped into regions based on altitude and geography. Points are colored according to their regional grouping for each species. *D. santomea* sites were pooled for all analyses. Topographic contour lines delineate altitude along the Pico de São Tomé transect.

Wild-caught females were placed individually into vials and allowed to lay eggs in the field on instant media (Nutri-Fly Instant Formulation, Genesee Scientific) supplemented with active yeast. This enabled us to establish isofemale lines and to sample F1 offspring that we preserved in ethanol upon emergence in the field. We used PCR to assess the infection status of each line from each site. The proportion of infected isofemale lines served as our estimate of *p* for each site, region, and species. To estimate *μ*, we assayed the *Wolbachia* infection status of newly emerged F1 offspring and then determined the proportion of uninfected adults produced by each wild-caught *Wolbachia*-infected female in the field. Importantly, because we sampled infected females from nature, the infection status of the male in these crosses is unknown. In crosses with infected males, uninfected ova produced by infected females could be lost if they are susceptible to CI (see Equation 2 from Turelli and Hoffmann [1995] for further dissection of this point). All PCR assays used primers for the *Wolbachia* surface protein (*wsp*), and a second set of primers for the arthropod-specific *28S* rDNA, which served as a positive control (Cooper *et al.* 2017; Meany *et al.* 2019). There is no evidence that *D. yakuba*-clade hosts are infected with other unaccounted for heritable symbionts, such as *Spiroplasma* (Mateos *et al.* 2006).

All statistical analyses were performed in R (R Core Team 2018). For each species, we first generated pooled estimates of *p* and *μ* that included all 2018 trapping sites. We then examined variation in *p* and *μ* by region and by trapping site. Assuming a binomial distribution, we estimated exact 95% binomial confidence intervals for *p* for each species/region/trapping site using the “binconf” function in the package *Hmisc* (Harrell and Dupont 2018). We used Fisher’s exact tests to assess pairwise differences in *p* among species and among regions. Estimates of *p* from 2018 were also compared to previously published data on São Tomé from Cooper *et al.* (2017); however, these prior estimates lacked adequate sampling to examine regional variation in *w*Yak and *w*San frequencies.

We estimated *μ* for each species/region/trapping site as the weighted mean of uninfected offspring produced by infected mothers, across all infected families, weighted by the number of F1s in each family. We then estimated 95% bias-corrected and accelerated bootstrap (BC_a_) confidence intervals using the “boot.ci” function and 5000 iterations in the *boot* package in R (Canty and Ripley 2017). BC_a_ confidence intervals are calculated using the two-sample acceleration constant given by equation 15.36 of Efron and Tibshirani (1993). Prior work has demonstrated substantial heterogeneity in *μ* among wild-caught *w*Ri-infected *D. simulans* (Turelli and Hoffmann 1995; Carrington *et al.* 2011) and *w*Suz-infected *D. suzukii* females (Hamm *et al.* 2014). Therefore, we visualized the full distribution of *μ* values for *w*Yak and *w*San, and used Kruskal-Wallis tests to assess heterogeneity in *μ* among wild-caught females for each species (Fox and Weisberg 2011). We used Kruskal-Wallis tests to assess differences in *μ* among trapping sites and Wilcoxon rank sum tests to assess pairwise differences in *μ* between species and between regions. Prior work also suggests maternal transmission of *Wolbachia* infecting *D. pseudotakahashii* is sex-biased, with perfect transmission to female progeny and reduced transmission to males (Richardson *et al.* 2018). Thus, we also evaluated whether imperfect transmission varied between female and male progeny for *w*Yak or *w*San.

### Determining environmental effects on imperfect *w* Yak maternal transmission

Our 2018 sampling revealed that transmission rates vary spatially on São Tomé for *w*Yak but not *w*San (see Results below). Specifically, *μ* was significantly higher for *w*Yak at high altitude, where conditions are relatively cool compared to low altitude. *Wolbachia* that are imperfectly transmitted in the field are perfectly or near perfectly transmitted under standard laboratory conditions (Hoffmann *et al.* 1990; Meany *et al.* 2019), suggesting that abiotic conditions may contribute to imperfect transmission in nature. Moreover, Turelli and Hoffmann (1995) found evidence that *w*Ri maternal transmission becomes near perfect when isofemale lines of *D. simulans* are cultured in the lab for six months. Environmental conditions can also influence other traits like cellular *Wolbachia* titer and localization in developing host oocytes, both of which are predicted to influence *μ* (Clancy and Hoffmann 1998; Serbus and Sullivan 2007; Ross *et al.* 2017; Christensen *et al.* 2019; Ross *et al.* 2019a). Thus, we manipulated environmental conditions in the laboratory in an attempt to determine the basis of variable imperfect *w*Yak transmission on São Tomé.

We first tested whether changes to the standard fly food diet alter *w*Yak transmission rates. Female *D. melanogaster* reared on a yeast-enriched diet for two days have reduced cellular *w*Mel titer in developing stage 10a oocytes (Serbus *et al.* 2015; Christensen *et al.* 2019). Reductions in oocyte titer are predicted to generate imperfect transmission, although this hypothesis has not been tested. We reared an infected *D. yakuba* isofemale line (*L5*; Table S2) at 25°C on a standard food diet and then placed newly emerged virgin females individually with two males of the same infected genotype in vials containing yeast-enriched food. Yeast-enriched food was prepared by mixing 1.5 ml of heat-killed yeast paste into 3.5 ml of standard food. Each subline was allowed to lay eggs for one week, and in the following generation, newly emerged male and female F1 offspring were screened for *w*Yak infection individually using PCR as described above.

Because imperfect maternal transmission was greatest at high altitude where conditions are generally cool (Table S1), we next tested whether rearing *D. yakuba* in cold conditions (20°C) perturbs maternal transmission relative to standard laboratory conditions (25°C). World Bioclim data (www.worldclim.org) indicate that 20°C is well within the average temperature range of our trapping sites on São Tomé (Table S1), and previous work has shown that *D. yakuba* experience reduced larval survival, egg hatch, and longevity at temperatures <20°C (Matute *et al.* 2009; Cooper *et al.* 2018). Thus, we reared the *L5* isofemale line in a 20°C incubator under a 12L:12D light cycle (Pericival Model I-36LL). Virgin females were placed individually with two males into vials and allowed to lay eggs for one week. Newly emerged male and female F1s from each subline were screened for *w*Yak infection using PCR as described above.

The 20°C cold treatment generated a significant increase in imperfect maternal transmission, whereas yeast-enriched food did not (see Results). We therefore investigated whether rearing *L5* females at 20°C reduces *w*Yak titer, a trait predicted to determine *Wolbachia* transmission. We used qPCR to compare titer in females reared at 20°C to those reared under standard conditions at 25°C. Females from each temperature treatment were aged to 3 days and then homogenized together in groups of 10. The final sample included 10 biological replicates of each temperature treatment. DNA was extracted using a DNeasy Blood & Tissue Kit (Qiagen). We used a Stratagene Mx3000P (Agilent Technologies) and qPCR primers designed for *D. yakuba* to amplify a *Drosophila*-specific locus (*Rpl32*; F: 5’-CCGCTTCAAGGGACAGTATC-3’, R: 5’-CGATCTCCTTGCGCTTCTTG-3’) and a *Wolbachia*-specific locus (*ftsZ*; F: 5’-ATCCTTAACTGCGGCTCTTG-3’, R: 5’-TTCATCACAGCAGGAATGGG-3’) with the following cycling conditions: 50°C for 2 minutes, 95°C for 2 minutes, and then 40 cycles of 95°C for 15 seconds, 58°C for 15 seconds, and 72°C for 1 minute. Efficiency curves were generated to confirm that primer efficiency was between 90–100% for *Rpl32* (96.58%) and *ftsZ* (93.24%). We used the average cycle threshold (*Ct*) value of three technical replicates for each sample. We estimated relative *Wolbachia* density as 2^Δ*Ct*^, where Δ*Ct* = *Ct*_*Rpl32*_ − *Ct*_*ftsZ*_ (Pfaffl 2001). We then used a Wilcoxon rank sum test to assess differences in titer between females reared at 20° and 25°C.

### Dissecting *Wolbachia* and host effects on *w*Yak maternal transmission

We next investigated the contributions of *w*Yak and *D. yakuba* genomes to imperfect *w*Yak transmission. Prior work indicates very little differentiation among *w*Yak genomes from West Africa (0.0007% third-position pairwise differences; Cooper *et al.* 2019), suggesting *w*Yak genomic variation may have little influence on variation in imperfect *w*Yak transmission. In contrast, host genomes have been shown to influence *Wolbachia* titer in developing host oocytes, which is predicted to determine transmission fidelity (Serbus and Sullivan 2007; Funkhouser-Jones *et al.* 2018). The genomes of *D. yakuba* sampled from low and high altitudes on São Tomé are weakly, but significantly, differentiated (F_ST_ = 0.0503, *P* < 0.001; Comeault *et al.* 2016; Turissini and Matute 2017). While we are ignorant of the type of host factors that modify *Wolbachia* titer and transmission in *Drosophila*, we characterized allele frequency and fixed differences between *D. yakuba* at low and high altitudes. In addition to this genomic characterization, we explicitly tested for host and *Wolbachia* contributions to variation in *μ* using genotypes with reciprocally introgressed *D. yakuba* and *w*Yak genomes.

Genomic reads were obtained from the published data of Turissini and Matute (2017) for *D. yakuba* from low (*N* = 7) and high altitude (*N* = 22) on São Tomé. As described in Turissini and Matute (2017), reads were mapped to version 1.04 of the *D. yakuba* reference genome (Drosophila 12 Genomes Consortium 2007) using bwa version 0.7.12 (Li and Durbin 2010). We merged BAM files with SAMtools version 0.1.19 (Li *et al.* 2009). Reads were then remapped locally in the merged BAM files using GATK version 3.2-2 using the RealignerTargetCreator and IndelRealigner functions (McKenna *et al.* 2010; DePristo *et al.* 2011). We used the GATK UnifiedGenotyper with the parameter het = 0.01, and applied QD = 2.0, FS_filter = 60.0, MQ_filter = 30.0, MQ_Rank_Sum_filter = −8.0 to the resulting vcf file. Sites were excluded if the coverage was less than 5 or the coverage was greater than the 99th quantile of the distribution of genomic coverage for each line or if the SNP failed to pass any of the GATK filters. All polymorphic sites from the resulting vcf file were stored in a MySQL table for analyses.

For each SNP, we calculated the difference in allele frequencies between low vs. high altitude *D. yakuba* on São Tomé. We summarized mean allelic differences along each chromosome using a sliding widow with 5kb increments. We calculated the mean and standard deviation of the full distribution of allele frequency differences using the function “tapply” in R. We also identified fixed differences across the genome. Finally, we tested whether the mean allele frequency difference observed along the genome differed from zero using a parametric Welch test and a nonparametric Wilcoxon rank sum test, which both agreed. We generated a normal distribution with 6,528,464 observations (the number of polymorphic sites) using the R function “rnorm” matching the same standard deviation observed in the empirical genome-wide data but centered on zero. We used the function “compare.2.vectors” in the *afex* package (Singmann *et al.* 2016) to compare the observed and simulated allele frequency distributions.

To explicitly dissect *w*Yak and *D. yakuba* contributions to variation in *μ*, we reciprocally introgressed *w*Yak and *D. yakuba* genomes using two infected *D. yakuba* isofemale lines as starting material (*L42* and *L48*; Table S2). We first crossed *L42* females with *L48* males and then backcrossed F1 females to *L48* males. We repeated this cross for five generations to generate the *L48*^*L42*^ genotype (*w*Yak variant denoted by superscript) composed of ~97% of the *L48* nuclear background and the introgressed *L42* cytoplasm. We then generated the reciprocal genotype (*L42*^*L48*^) using the same approach. We measured *μ* for the two naturally sampled genotypes (*L42* and *L48*) and for the two reciprocally introgressed genotypes (*L42*^*L48*^ and *L48*^*L42*^) reared under standard laboratory conditions (25°C, 12L:12D). Virgin females were placed individually with two males of the same genotype into vials and allowed to lay eggs for one week. Newly emerged male and female F1 offspring were screened for *w*Yak infection individually using PCR (as described above). We first tested whether *μ* varied among the four genotypes (*L42*, *L48*, *L42*^*L48*^, *L48*^*L42*^) using a Kruskal-Wallis test. Because *μ* takes the form of a frequency, we also used a generalized linear model (GLM) to assess the contributions of the host and *Wolbachia* genomes to imperfect transmission. We used the “glm” function in R to fit a GLM with a Poisson error structure and used the raw count data of uninfected F1s from each family as the dependent variable. We then included the host nuclear genome, the *Wolbachia* genome, and their interaction as independent variables. Both the Kruskal-Wallis test and the GLM revealed no evidence for host or *Wolbachia* effects on *μ*, so only the Kruskal-Wallis results are presented herein.

### Modeling infection equilibria with field estimates of *p* and *μ*

To explore the relationship between our field estimates of *p* and *μ* on São Tomé, we considered an idealized discrete-generation model for *Wolbachia* frequency dynamics proposed by Hoffmann et al. (1990). This model incorporates *μ*, *F* (*Wolbachia* effects on host fitness), and *H* (the severity of CI) (Hoffmann and Turelli 1997)—we previously estimated the latter two parameters in the lab (Cooper *et al.* 2017). In cases of imperfect transmission, the model assumes that uninfected ova are equally susceptible to CI regardless of the infection status of their mother, which is supported by results from *w*Ri-infected *D. simulans* (Turelli and Hoffmann 1995; Carrington *et al.* 2011). Embryos produced by uninfected mothers mated with infected fathers hatch with frequency *H* = 1 – *s*_*h*_, relative to the other three possible fertilizations, which are all considered equally compatible (Cooper *et al.* 2017). Thus, *s*_*h*_ represents the severity of CI, or the frequency of unhatched eggs in pairings between uninfected females and infected males.

Prior estimates of CI indicate that *w*Yak and *w*San reduce egg-to-adult viability of uninfected females mated to infected males by about 10 to 15%, relative to compatible crosses (Cooper *et al.* 2017). We first ignored this weak CI and considered a stable equilibrium 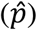 balanced by imperfect transmission (*μ* > 0) and positive fitness effects (*F* > 1). Benefits to host fitness have yet to be directly connected to low frequency *Wolbachia* spread in these systems or any others, but like Hoffmann and Turelli (1997) conjectured for non-CI-causing *Wolbachia*, we assume *F*(1 – *μ*) must be greater than 1 given the spread and persistence of *w*Yak and *w*San in nature (Cooper *et al.* 2017). When *F*(1 – *μ*) > 1, the stable equilibrium frequency is

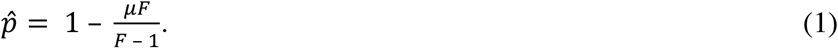

In this case, 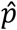 increases from 0 toward 1 – *μ* as *F* increases from 1/(1 – *μ*). We incorporated our field estimates of imperfect maternal transmission to explore how *μ* and a range of positive *Wolbachia* fitness effects might explain observed estimates of *p* for *w*Yak and *w*San.

Next, we considered equilibria that incorporate CI (i.e., *s*_*h*_ > 0) (Turelli and Hoffmann 1995; Kriesner *et al.* 2016). CI does not contribute to the initial spread of *Wolbachia*, yet strong-CI-causing strains like *w*Ri spread rapidly to high equilibrium frequencies (Turelli and Hoffmann 1995; Kriesner *et al.* 2013). Thus, we assume CI-causing strains must also increase host fitness to initially spread from low frequency such that *F*(1 – *μ*) > 1. CI-causing infections that generate *F*(1 – *μ*) > 1 and *Fμ* < 1 produce a single stable equilibrium between 0 and 1 given by

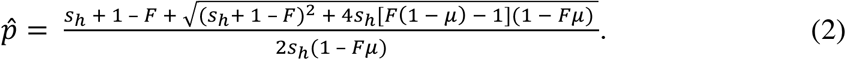

Laboratory estimates of CI in the *D. yakuba* clade support *s*_*h*_ < 0.2 for both *w*Yak (*s*_*h*_ = 0.16) and *w*San (*s*_*h*_ = 0.15) (Cooper *et al.* 2017). In contrast, the relatively high infection frequencies of *w*Yak and *w*San in 2018 (Figure 2, Table 1) are a hallmark of strong CI (e.g., Turelli and Hoffmann 1995), implying these *Wolbachia* may cause stronger CI in the field. Thus, we considered stronger CI (*s*_*h*_ = 0.20 and *s*_*h*_ = 0.45), in addition to our laboratory estimates of *s*_*h*_ for each *Wolbachia*.

**FIGURE 2.**
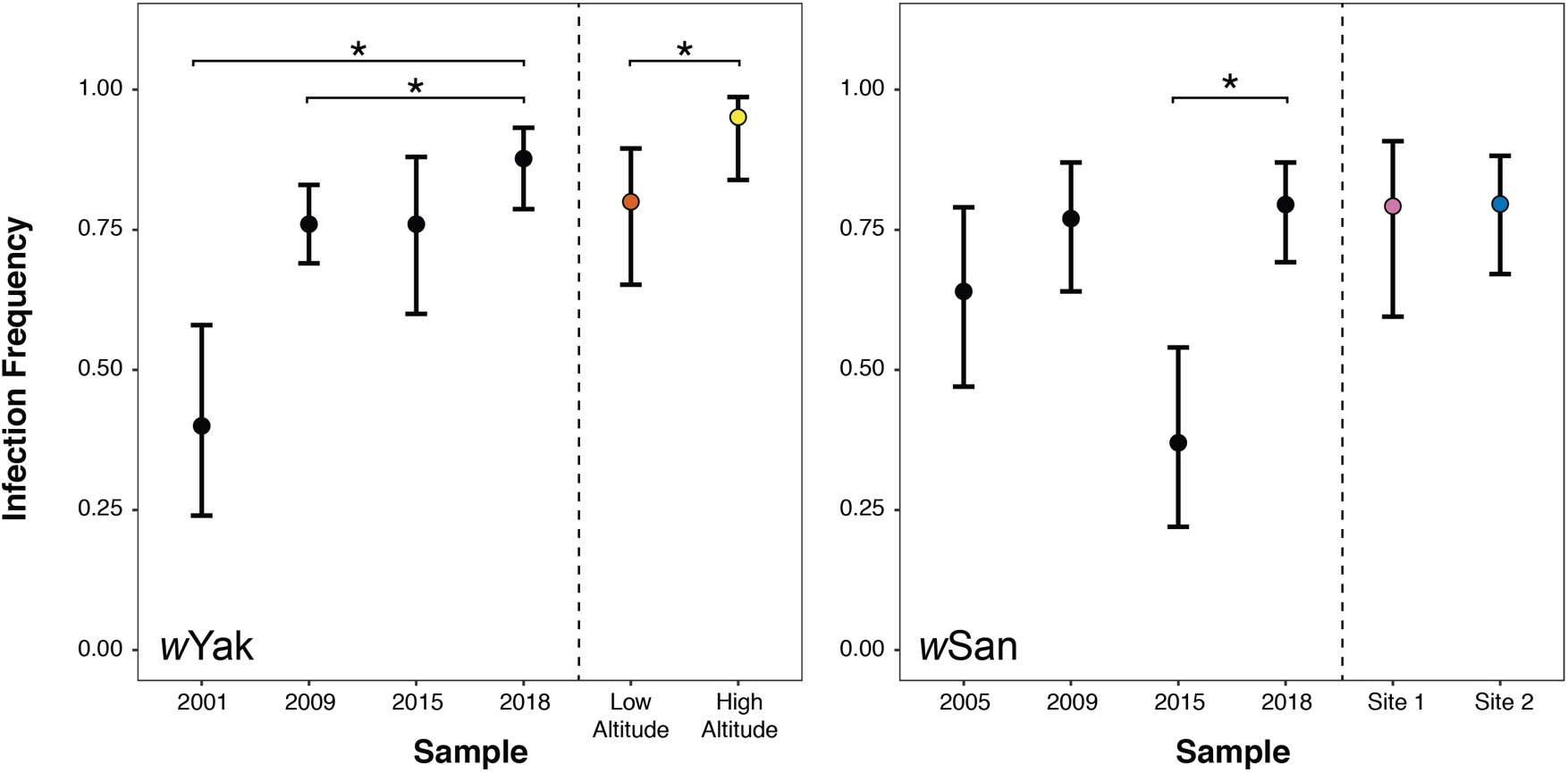
*Wolbachia* infection frequencies on São Tomé. Data from the present study (2018) are compared to previous estimates of infection frequencies from 2001-2015 (Cooper *et al.* 2017). Error bars represent 95% binomial confidence intervals. To the right of vertical dashed lines, 2018 infection frequencies are separated by region for each species. Points are color-coded according to Figure 1. Asterisks indicate statistically significant differences between 2018 and prior years, or between regions in the 2018 sample at *P* < 0.05.

**TABLE 1.** *Wolbachia* infection frequencies in *D. yakuba* and *D. santomea* on São Tomé.

| Species | Year | *N* | Infected | *p* [Confidence Interval] |
| --- | --- | --- | --- | --- |
| *D. yakuba* | 2001 | 35 | 14 | 0.40 [0.24, 0.58] |
|  | 2009 | 155 | 118 | 0.76 [0.69, 0.83] |
|  | 2015 | 41 | 31 | 0.76 [0.60, 0.88] |
|  | 2018 | 81 | 71 | 0.88 [0.79, 0.93] |
|  | Low altitude | 40 | 32 | 0.80 [0.65, 0.90] |
|  | High altitude | 41 | 39 | 0.95 [0.84, 0.99] |
| *D. santomea* | 2005 | 39 | 25 | 0.64 [0.47, 0.79] |
|  | 2009 | 57 | 44 | 0.77 [0.64, 0.87] |
|  | 2015 | 38 | 14 | 0.37 [0.22, 0.54] |
|  | 2018 | 78 | 62 | 0.80 [0.69, 0.87] |
|  | Site 1 | 24 | 19 | 0.79 [0.60, 0.91] |
|  | Site 2 | 54 | 43 | 0.80 [0.67, 0.88] |

Sample sizes (*N*), infection frequencies (*p*), and exact 95% binomial confidence intervals are shown for each year. Data from 2001-2015 are from Cooper *et al.* (2017). For each species, infection frequencies are calculated for the pooled 2018 dataset and for each region.

Our analysis of *w*Yak infection equilibria at high altitude revealed that biologically unrealistic parameter values (e.g., *F* > 4, *s*_*h*_ > 0.45) are required to explain the observed combination of high *w*Yak frequency (*p* = 0.95 [0.84, 0.99]) and very imperfect *w*Yak transmission (*μ* = 0.20 [0.087, 0.364]). We explored two additional processes the might contribute to this counterintuitive pattern at high altitude: (1) temporal variation in imperfect maternal transmission and (2) stochastic infection frequency fluctuations.

First, we consider temporal variation in *μ*. Discrete generation models use estimates of *μ*, *F*, and *H* to infer 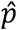 in the next host generation. We estimated *μ* and *p* concurrently in the field, assuming that our estimates of *μ* in the current generation reflect *μ* in the prior host generation. Turelli and Hoffmann (1995) found that imperfect *w*Ri transmission decreased significantly from 0.044 [0.029, 0.090] in April to 0 [0.000, 0.007] in November of 1993 (*P* < 0.001) in an Ivanhoe, California population of *D. simulans* (Carrington *et al.* 2011), but it remains unknown whether *Wolbachia* transmission rates vary over shorter timescales within the same host population (e.g., across a single host generation). We repeated our mathematical analyses to consider the full range of *μ* point estimates across all trapping sites on São Tomé (Table S1). This range exceeds the seasonal variation in *μ* estimated for *w*Ri in Ivanhoe, CA, and we reasoned that any generation-to-generation variation in imperfect transmission within a single location is likely less than the range of point estimates across all trapping sites. Thus, this analysis conservatively considers how different values of *μ* in the prior host generation would alter 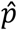.

Next, we consider the potential for stochastic *w*Yak frequency fluctuations to influence the patterns we observed. We explored the effect of host population size on *w*Yak frequency fluctuations by modifying the model of Hoffmann and Turelli (1997) as described in Kreisner and Hoffmann (2018) to incorporate randomly variable outcomes for *μ*, *F*, and *H*, as well as for male mating success, crossing types arising from matings, female reproductive success, and the chance of viable embryos surviving to adulthood. We selected plausible combinations of *μ*, *F*, and *H* for *w*Yak at low and high altitude (based on the abovementioned mathematical analysis; Table S5) to illustrate how stochasticity contributes to variation in *w*Yak frequencies at 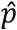. Monte Carlo simulations with 10,000 replicates of population events were enacted using functions in the PopTools package (Hood 2011) and the parameters described above (see Supplemental Methods for a detailed description). The initial generation for each replicate trial comprised a set number of adults with two infection status types: uninfected and *w*Yak-infected. Each infection type consisted of an equal number of females and males, and we assumed females had pre-mated with a male of the same infection type. Trials were initiated with the host population at 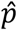 and were run for 100 host generations. To evaluate how host population size affects stochasticity, we tested three different host census population sizes (*N* = 5,000, 50,000, and 500,000). Every 20 host generations, we calculated the mean infection frequency of the Monte Carlo simulations and associated 95% confidence intervals.

### Data Accessibility

All supplemental materials, data, and scripts will be made available on FigShare. Genomic data from Turissini and Matute (2017) are available at NCBI (Bioproject: PRJNA395473).

## RESULTS

### *w*Yak and *w*San infection frequencies vary on São Tomé

*w*Yak and *w*San frequencies vary through time on São Tomé, and *w*Yak frequencies vary spatially within São Tomé and between São Tomé and Bioko (Figure 2, Table 1, Cooper *et al.* 2017). Across all *D. yakuba* trapping sites, our pooled estimate of *w*Yak frequency in 2018 (*p* = 0.88) is significantly higher than our estimates from 2001 (*p* = 0.40; Fisher’s Exact *P* < 0.0001) and 2009 (*p* = 0.76; Fisher’s Exact *P* = 0.040), although 2018 did not differ from our 2015 estimate (*p* = 0.76; Fisher’s Exact *P* = 0.120) (Cooper *et al.* 2017). In 2018, *w*Yak frequencies varied regionally, such that *w*Yak frequency was significantly higher at high altitude (*p* = 0.95; yellow sites in Figure 1) than at low altitude (*p* = 0.80; orange sites in Figure 1) along the Pico de São Tomé transect (Fisher’s Exact *P* = 0.048). While this is the first demonstration of within island spatial (altitudinal) variation in *p*, it is consistent with previous observations of geographic variation in *w*Yak frequencies in 2009 between São Tomé (*p* = 0.76) and the neighboring island of Bioko (*p* = 0.03; *P* < 0.001) (Cooper *et al.* 2017).

Across all *D. santomea* trapping sites, the pooled frequency of *w*San in 2018 was relatively high (*p* = 0.80), and did not differ statistically from past estimates on the island in 2005 (*p* = 0.64; Fisher’s Exact *P* = 0.115) and 2009 (*p* = 0.77; Fisher’s Exact *P* = 0.833). The 2018 estimate was significantly greater than the most recent 2015 estimate (*p* = 0.37; Fisher’s Exact *P* < 0.0001), providing further support for temporal variation in *w*San frequencies on São Tomé (Cooper *et al.* 2017). *w*San frequencies varied between years when *w*Yak frequencies were relatively stable, and vice versa, suggesting these nearly identical *Wolbachia* behave differently across host species backgrounds and/or abiotic environments. Unlike *w*Yak, we did not find evidence for regional variation in *w*San frequencies, such that site 1 (*p* = 0.79; purple sites in Figure 1) and site 2 (*p* = 0.80; blue sites in Figure 1) were statistically indistinguishable (Fisher’s Exact *P* = 1).

### Imperfect maternal transmission varies spatially for *w*Yak, but not *w*San

Imperfect maternal transmission was heterogeneous among wild-caught *D. yakuba* and *D. santomea* females. *Wolbachia* were usually transmitted with high fidelity, but some females exhibited substantial imperfect transmission (Figure 3). Accordingly, we found significant heterogeneity in *μ* among wild-caught females for *D. yakuba* (*χ*^2^ = 579.2, *P* < 0.001) and *D. santomea* (*χ*^2^ = 418.8, *P* < 0.001). Pooled *μ* values across all trapping sites (Table 2) did not differ between *w*Yak (*μ* = 0.126) and *w*San (*μ* = 0.068; *W* = 2355.5, *P* = 0.283). Contrary to our predictions, regional estimates of *μ* for *w*Yak and *w*San were not negatively correlated with *p* (*F*_1,2_ = 5.333, *P* = 0.147). This remained true when families were grouped by individual trapping sites (e.g., YAK02, CAR01, etc.) rather than region (*F*_1,15_ = 0.078, *P* = 0.784).

**FIGURE 3.**
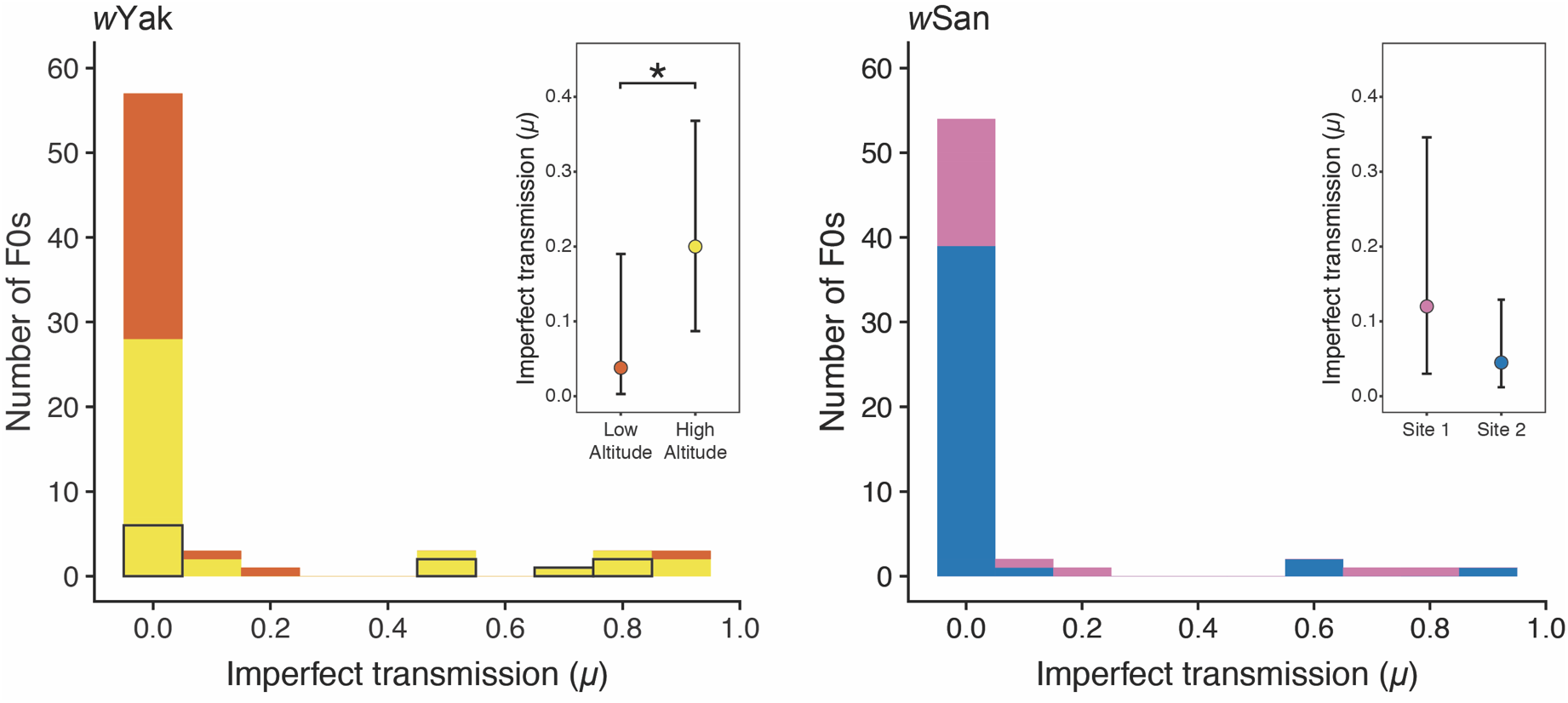
Histogram of *μ* for *w*Yak-infected *D. yakuba* (left) and *w*San-infected *D. santomea* (right) females. The bars are color coded by region as in Figure 1. For *w*Yak, the contributions of F0 females from the anomalous site YAK05b are outlined in black (see main text). Insets show mean estimates of *μ* and associated 95% BC_a_ confidence intervals for each region. Asterisk indicates a statistically significant difference in *w*Yak *μ* between low and high altitude at *P* < 0.05.

**TABLE 2.** Imperfect maternal *Wolbachia* transmission by *D. yakuba* and *D. santomea* on São Tomé.

| Species | Year | Mean *N* F1s | $\mu$ [Confidence Interval] |
| --- | --- | --- | --- |
| *D. yakuba* | 2018 | 10.05 | 0.126 [0.062, 0.238] |
|  | Low Altitude | 9.85 | 0.038 [0.003, 0.184] |
|  | High Altitude | 10.24 | 0.20 [0.087, 0.364] |
| *D. santomea* | 2018 | 9.80 | 0.068 [0.027, 0.154] |
|  | Site 1 | 9.96 | 0.120 [0.032, 0.346] |
|  | Site 2 | 8.64 | 0.045 [0.011, 0.130] |

Mean number of F1 offspring per family (Mean *N* F1s), weighted mean imperfect transmission ( $\mu$ ), and 95% BC<sub>a</sub> confidence intervals are shown for 2018. For each species, rates of maternal transmission are calculated for the pooled 2018 dataset and for each region.

Differences in *w*Yak *μ* among individual trapping sites were not statistically significant (*χ*^2^ = 7.6, *P* = 0.055; Table S1); however, when traps where grouped into altitudinal regions, we detected regional variation in *w*Yak *μ*, such that maternal transmission was more imperfect at high altitude (*μ* = 0.20) relative to low altitude (*μ* = 0.038; *W* = 503, *P* = 0.045; Table 2, Figure 3). This is surprising given that *w*Yak frequencies were significantly higher at high altitude compared to low altitude. To explain this counterintuitive relationship between *w*Yak *μ* and *p*, either *μ* must vary on short timescales within regions or *F* and/or *H* must vary between regions (see below). Finally, we found no evidence for differences in *w*Yak transmission to male (*μ* = 0.192 [0.087, 0.346]) and female (*μ* = 0.204 [0.084, 0.386]) offspring at high altitude (*W* = 722.5, *P* = 0.994), or to males (*μ* = 0.033 [0, 0.172]) and females (*μ* = 0.043 [0, 0.216]) at low altitude (*W* = 512.5, *P* = 0.551).

Imperfect *w*San transmission did not vary among trapping sites (*χ*^2^ = 16.0, *P* = 0.098; Table S1) or between regional groupings, such that site 1 (*μ* = 0.120) and site 2 (*μ* = 0.045) were statistically indistinguishable (*W* = 456.5, *P* = 0.213; Table 2, Figure 3). This is not surprising given only a mountainous ridge separates these sites (Figure 1). As with *w*Yak, we observed no difference between rates of *w*San transmission to male (*μ* = 0.129 [0.027, 0.341]) and female (*μ* = 0.114 [0.03, 0.378]) offspring at site 1 (*W* = 190, *P* = 0.698) or to male (*μ* = 0.054 [0.013, 0.144]) and female (*μ* = 0.038 [0.004, 0.127]) offspring at site 2 (*W* = 866, *P* = 0.779). Because we found no evidence for variation in *μ* or *p* for *w*San, we disregarded regional groupings and used the pooled 2018 parameter estimates (*μ* = 0.068, *p* = 0.795) when modeling infection equilibria (see below). The pooled estimate of *μ* = 0.068 for *w*San was statistically indistinguishable from *w*Yak at low (*W* = 1028.5, *P* = 0.607) and at high altitude (*W* = 1018, *P* = 0.0513).

### Cool rearing temperature increases imperfect maternal transmission

In the laboratory, maternal transmission was near perfect when infected-females were placed on yeast-enriched food (*μ* = 0.011; Figure 4, Table S3). *μ* values from the yeast treatment were homogenous among individual mothers (*χ*^2^ = 25.9, *P* = 0.523; Figure S1). Compared to our field data, *μ* on yeast-enriched food was not significantly different than *μ* at low altitude (*W* = 440, *P* = 0.557), but was significantly lower than *μ* at high altitude (*W* = 440, *P* = 0.030) (Figures 3 and 4). Imperfect transmission on yeast-enriched food also did not vary between males (*μ* = 0) and females (*μ* = 0.021 [0.007, 0.058]; *W* = 434, *P* = 0.081).

**FIGURE 4.**
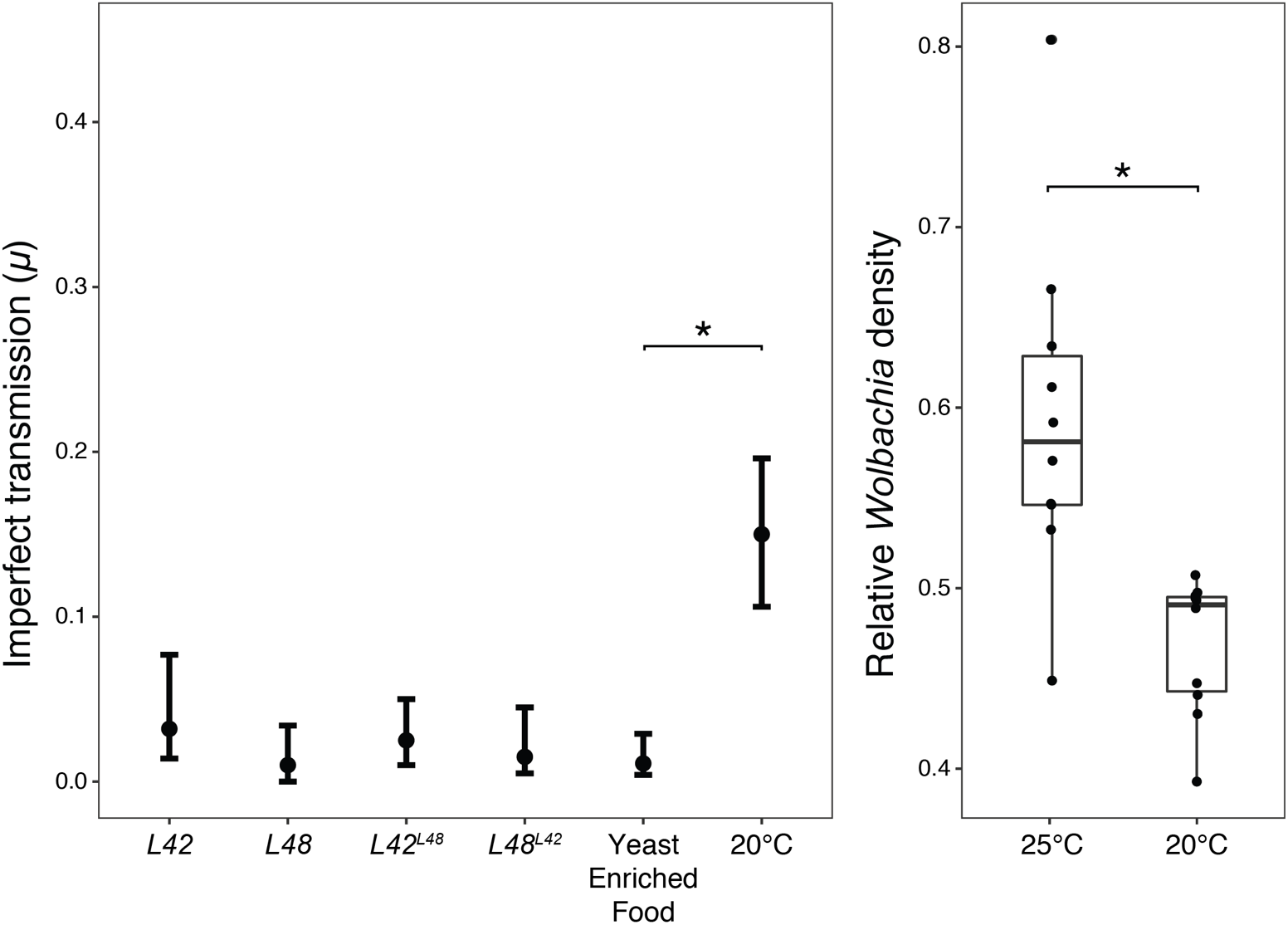
Laboratory estimates of imperfect *w*Yak transmission and titer. Left) Mean *μ* and 95% confidence intervals for *w*Yak from laboratory experiments dissecting genetic and environmental contributions to imperfect maternal transmission. *μ* values are shown for naturally sampled (*L42*, *L48*) and for reciprocally introgressed genotypes (*L42*^*L48*^, *L48*^*L42*^) reared under standard conditions (25°C), and for *L5* reared on yeast-enriched food at 25°C and on standard food at 20°C. Right) Estimates of *w*Yak titer (relative *Wolbachia* density) for *L5* females reared at 25° and 20°C. Asterisks indicate statistically significant differences at *P* < 0.05.

In contrast, we detected substantial imperfect *w*Yak transmission when infected *D. yakuba* females were reared at relatively cool 20°C (*μ* = 0.15; Figure 4, Table S3). *μ* values at 20°C were homogenous among individual mothers (*χ*^2^ = 16.5, *P* = 0.627; Figure S1). *μ* from the cold treatment was significantly greater than our field estimate at low altitude on São Tomé (*W* = 548, *P* < 0.001), but it was significantly lower than *μ* at high altitude (*W* = 535, *P* = 0.011) (Figures 3 and 4). *w*Yak transmission under cold exposure was also more imperfect than in the yeast-enriched food treatment (*W* = 74, *P* < 0.001; Figure 4). When reared at 20°C, *D. yakuba* females also had a lower relative *w*Yak density (2^Δ*Ct*^ = 0.469 [0.441, 0.496]) than females developed at 25°C (2^Δ*Ct*^ = 0.595 [0.527, 0.663]; *W* = 6, *P* < 0.001; Figure 4). We predict that reductions in *w*Yak titer at cool temperatures contribute to imperfect maternal transmission. Interestingly, maternal transmission to female progeny in our 20°C treatment was an order of magnitude more imperfect (*μ* = 0.277 [0.118, 0.376]) than to males (*μ* = 0.02 [0, 0.065]; *W* = 342, *P* < 0.001), a pattern opposite that observed for the *Wolbachia* infecting *D. pseudotakahashii* (Richardson *et al.* 2019). This pattern is also opposite our theoretical expectation that selection should favor faithful transmission to female offspring since their uninfected ova are susceptible to CI (Prout 1994; Turelli 1994). While transmission to females and males did not differ in the field on São Tomé or for our yeast-enriched food treatment, in all cases our point estimates for *w*Yak *μ* were higher for females than males.

### No evidence for *Wolbachia* or host effects on maternal transmission

We found very little differentiation in allele frequencies along the genomes of *D. yakuba* from low and high altitudes (Figure S2). The mean difference in allele frequencies between low and high altitude *D. yakuba* was 0.0027 (lower 25% quartile = 0, upper quartile = 0.0455), which is significantly higher than zero (Welch Two Sample t-test: t = 51.42, df = 13,056,636, *P* < 0.0001); however, the simulated distribution centered on 0 and our observed distribution are almost completely overlapping (Figure S2). This is consistent with two prior studies showing very low differentiation between these two regions (Comeault *et al.* 2016; Turissini and Matute 2017). Only eight of the 6,528,464 SNPs across the genome were fixed for different nucleotides between the low and high *D. yakuba* populations. None of these fixed differences were located in protein-coding sequence, but all eight SNPs occurred within 10kb of genes (Table S4). All nearby genes were either functionally undescribed or had no obvious connection to interactions with *Wolbachia.* We are ignorant of the types of genes that might modify *w*Yak transmission and have no reason to think these fixed differences between low and high altitude *D. yakuba* influence the variation in transmission we observe.

To explicitly test for host modulation of *w*Yak transmission, we reciprocally introgressed *w*Yak and *D. yakuba* genomes and quantified *μ* for the starting (*L42* and *L48*) and reciprocally introgressed (*L42*^*L48*^ and *L48*^*L42*^) genotypes. Maternal transmission was near perfect for all four genotypes when reared under standard 25°C conditions (Table S3). We found no evidence for *D. yakuba* effects on *w*Yak transmission, such that *μ* did not differ among the four genotypes (*χ*^2^ = 2.5, *P* = 0.478). For the *L42* genotype (*μ* = 0.032), *μ* was homogeneous among mothers (*χ*^2^ = 27.0, *P* = 0.171; Figure S1), and *w*Yak transmission to female progeny was significantly more imperfect (*μ* = 0.042 [0.016, 0.105]) than to males (*μ* = 0; *W* = 286, *P* = 0.041). For the *L48* genotype (*μ* = 0.01), *μ* was also homogeneous among mothers (*χ*^2^ = 18.8, *P* = 0.535), but transmission to female (*μ* = 0.009 [0, 0.049]) and male progeny (*μ* = 0.01 [0, 0.061]) did not differ (*W* = 220, *P* = 1). For the reciprocally introgressed genotype *L42*^*L48*^ (*μ* = 0.024), *μ* was homogeneous among mothers (*χ*^2^ = 15.2, *P* = 0.709), and transmission to female progeny (*μ* = 0.05 [0.02, 0.103]) was again significantly more imperfect than to males (*μ* = 0; *W* = 250, *P* = 0.020). Finally, for the reciprocally introgressed genotype *L48*^*L42*^ (*μ* = 0.015), *μ* values were homogeneous among mothers (*χ*^2^ = 17.0, *P* = 0.523), but transmission to female (*μ* = 0.01 [0, 0.059]) and male progeny (*μ* = 0.01 [0, 0.061]) did not differ (*W* = 200, *P* = 1). Interestingly, we detected female-biased imperfect transmission only for genotypes with the *L42* host nuclear genome (*L42* and *L42*^*L48*^), motivating future analysis of *D. yakuba* factors with sex-specific effects on *w*Yak transmission.

### Temporally variable *μ* or spatially variable *F* and/or *H* is required to explain *w*Yak infection equilibria

We modeled 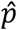, incorporating our field estimates of *μ* on São Tomé and lab estimates of *F* and *H* (Cooper *et al.* 2017). Beneficial *Wolbachia* effects of *F* > ~1.2 have not been documented in any system (Weeks *et al.* 2007; Meany *et al.* 2019), and prior efforts found no evidence for positive *Wolbachia* effects on *D. yakuba*-clade host fecundity (Cooper *et al.* 2017). *Wolbachia* effects on other components of host fitness have not been examined, but we assume *F*(1 – *μ*) must be greater than 1 given the spread and persistence of *w*Yak and *w*San in nature (Cooper *et al.* 2017). We generally consider values of *F* > 1.5 as biologically unrealistic (Meany *et al.* 2019). Finally, while *w*Yak and *w*San cause weak CI on average in the laboratory (*s*_*h*_ < 0.20), we considered equilibria ranging from no CI (*s*_*h*_ = 0) up to very strong CI (*s*_*h*_ = 0.45). Laboratory crosses and field infection frequencies suggest CI strength may vary in this clade (Cooper *et al.* 2017).

The results of our mathematical analyses are summarized in Figure 5. We conservatively evaluated infection equilibria across the full credible intervals for estimates of *μ* for each region/species (Figure S3, Table S5). Because *μ* could vary within sites across host generations, we also evaluated equilibria across the full range of *μ* point estimates for all trapping sites on São Tomé (Table S6). We reasoned that temporal variation in *μ* within any one site is likely lower than the full range of *μ* point estimates for all trapping sites on São Tomé, but future analysis of short-term variation in *μ* within field sites is required to test this assumption. Finally, we evaluated how host population size affects stochastic fluctuations in *Wolbachia* frequencies in host populations at infection equilibrium (Figure S4, Table S7).

**FIGURE 5.**
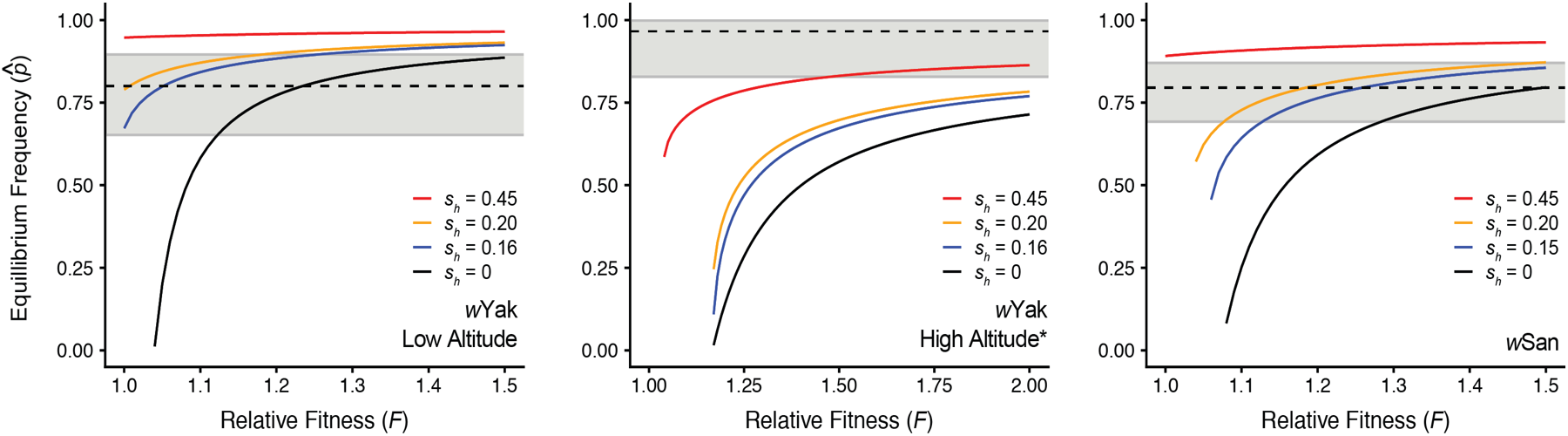
Equilibrium *w*Yak and *w*San infection frequencies plotted against a range of *F* values, assuming field estimates of *μ*. Dotted lines indicate observed infection frequencies for each region, and the gray area denotes 95% binomial confidence intervals for observed frequencies. Black lines denote no CI (*s*_*h*_ = 0), followed by laboratory estimates of weak CI in blue for *w*Yak (*s*_*h*_ = 0.16) and *w*San (*s*_*h*_ = 0.15) (Cooper *et al.* 2017), moderate CI in orange (*s*_*h*_ = 0.20), and strong CI in red (*s*_*h*_ = 0.45). The asterisk signifies that the YAK05b trapping site was removed from the high altitude *w*Yak dataset due to an anomalously high value of *μ* (see Table S1, Figure 3). Similar plots showing parameter estimates across the full credible intervals of *μ* are shown in Figure S3.

We first modeled infection equilibria for *w*Yak at low altitude. Given our estimate of *μ* = 0.038 [0.003, 0.184], no-to-moderate CI (*s*_*h*_ = 0-0.20), and modest host fitness effects (*F* = 1.01–1.23), we can plausibly explain our *w*Yak frequency estimate of *p* = 0.80 [0.65, 0.90] (Figure 5, Table S5). Assuming our lab estimate of weak CI (*s*_*h*_ = 0.16) and our point estimate of *μ* = 0.038, *F* = 1.25 is required to explain the upper credible interval for *p* (0.90) (Figure S3, Table S5). This suggests that our field estimate of imperfect transmission, combined with laboratory estimates of CI and modest-to-strong (positive) *Wolbachia* effects on host fitness, can reasonably explain the credible interval for *p* at low altitude. However, very unrealistic values of *F* combined with strong CI are almost always required to explain *p* if we assume the upper credible interval for *μ* at low altitude (0.184).

Next, we modeled infection equilibria for *w*Yak at high altitude, where we observed the counterintuitive pattern of a high infection frequency (*p* = 0.95 [0.84, 0.99]) despite very imperfect maternal transmission (*μ* = 0.20 [0.087, 0.364]). Using our point estimate for *μ*, stronger CI than previously estimated (*s*_*h*_ > 0.45) and biologically unrealistic positive fitness effects (*F* > 4) are required to explain *p* (Figure S3, Table S5; Cooper *et al.* 2017). Even considering the full credible interval for *μ*, unrealistic values of *F* > 1.5 are required to approach the credible interval of *p*, regardless of CI strength. If we assume *w*Yak transmission in the prior *D. yakuba* generation was substantially lower, such that *μ* = 0.038 (i.e., the lowest *w*Yak *μ* point estimate for all trapping sites on São Tomé), we can plausibly explain the lower credible limit of *p* (0.84) using our lab estimate of CI (*s*_*h*_ = 0.16) and *F* = 1.1 (Table S6).

Further exploration of our data indicates that imperfect transmission at one particular trapping site at high altitude (YAK05b, altitude = 1104 m) was more than three times higher than prior field estimates in any system (*μ* = 0.354; see Figure 3, Tables S1, S8). This is surprising given that the YAK05b site does not differ noticeably from the nearby trap YAK05, which is within eyesight of YAK05b (Figure 1). Excluding the anomalously high YAK05b site from the high altitude region, *p* = 0.97 [0.83, 0.998] (*N* = 29) and *μ* = 0.143 [0.036, 0.356]. Here, the lower credible interval of *p* can plausibly be explained by strong CI (*s*_*h*_ = 0.45) and *F* = 1.48 if we assume our point estimate of *μ* = 0.143, or by weak CI (*s*_*h*_ = 0.16) and *F* = 1.07 if we assume the lower credible interval of *μ* = 0.036 (Figure 5, Table S5). Similar values of *s*_*h*_ and *F* are required to explain *p* if we assume differences in imperfect maternal transmission in the prior host generation (Table S6).

We found no evidence that stochastic fluctuations in *w*Yak frequencies could influence the patterns we observe. We chose three different combinations of parameter estimates from Table S5 and evaluated how host population size affects stochasticity (Supplemental Methods, Figure S4, Table S7). In all cases, even small host populations are expected to yield deterministic infection dynamics. As an example, at high altitude our lower credible limit of *p* = 0.84 can be explained by assuming the lower credible limit of *μ* = 0.087, moderate CI of *s*_*h*_ = 0.2, and very strong fitness benefits of *F* = 1.55 (Table S5). Assuming an unrealistically small host population of 5,000 individuals and *w*Yak 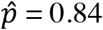, we inferred mean *p* = 0.84 [0.82, 0.86] after 100 simulated host generations (Figure S4, Table S7). When we removed the anomalous YAK05b site and assumed an infection equilibrium of 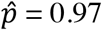 with *μ* = 0.036, *s*_*h*_ = 0.45, and *F* = 1.47 (Table S5), we found even less variance in infection frequencies with *p* = 0.966 [0.960, 0.972] after 100 generations. This is consistent with simulations and cage experiments using *w*Ri- and *w*Au-infected *D. simulans* showing that small experimental host populations tend to yield deterministic infection dynamics (Kreisner and Hoffmann 2018). These results imply that host population size fluctuations are unlikely to explain why concurrent estimates of *μ* do not predict *w*Yak infection frequencies at high altitude. Plausible explanations for *w*Yak at high altitude generally require us to infer (1) a higher fidelity of transmission in the *D. yakuba* generation prior to our concurrent estimation of *μ* and *p* and/or (2) more positive *w*Yak effects on *D. yakuba* fitness and/or stronger CI at high relative to low altitude. We predict that some combination of these two possibilities determines *w*Yak frequency variation on São Tomé.

We found no evidence for regional variation in *w*San *p* or *μ* on São Tomé, so we modeled infection equilibria for our pooled sample of *D. santomea*. Generally, we can explain *p* = 0.80 [0.69, 0.87] assuming our point estimate of *μ* = 0.068, weak-to-moderate CI (*s*_*h*_ = 0.15–0.2), and *F* < 1.5 (Figure 5, Table S5). For example, assuming *μ* = 0.068 and our laboratory estimate of CI strength (*s*_*h*_ = 0.15), *F* must equal 1.26 to explain *p* = 0.80. Assuming more imperfect *w*San transmission (e.g., *μ* = 0.154, the upper credible interval) we can still explain *p* = 0.80 with *F* < 1.5, but we must invoke strong CI (*s*_*h*_ = 0.45). Thus, unlike *w*Yak frequencies on São Tomé, we can plausibly explain much of our credible interval for *w*San *p* with our point estimate for *μ*, weak-to-moderate CI, and biologically reasonable values of *F*.

## DISCUSSION

*w*Mel-like *Wolbachia* in *Drosophila* tend to persist at intermediate frequencies that vary (Kriesner *et al.* 2016; Cooper *et al.* 2017), but the contributions of imperfect maternal transmission, *Wolbachia* fitness effects, and CI to frequency variation remain unknown. Our field analysis found that both *w*Yak and *w*San are imperfectly transmitted on São Tomé, and *w*Yak transmission varies spatially. Contrary to our prediction, concurrent field estimates of imperfect maternal transmission do not predict spatial variation in *w*Yak frequencies, which are highest at high altitudes where maternal transmission is the most imperfect. Genomic and genetic analyses suggest little contribution of host genomes to variation in *w*Yak maternal transmission. Instead, rearing *D. yakuba* females at a cool temperature significantly decreases *w*Yak titer and increases imperfect transmission to levels observed in nature. Using mathematical models, we infer that temporal variation in *w*Yak transmission within sites, and/or spatial variation in *w*Yak effects on host fitness and CI among sites, is required to explain *w*Yak frequencies. In contrast, while *w*San frequencies continue to vary across years (Cooper *et al.* 2017), they were spatially stable within São Tomé in 2018, and plausibly explained by imperfect transmission, modest fitness effects, and weak CI. We discuss the implications of our findings below.

### Environmental effects on maternal*Wolbachia* transmission

Altitudinal differences in *w*Yak frequency (*p* = 0.80 to 0.95) and imperfect maternal transmission (*μ* = 0.038 to 0.20) occur over short geographic distances, with low and high altitude *D. yakuba* sites separated by only 2.4 km and an elevation gain of 310 m (Table S1). Conditions are relatively cool at high altitude on Pico de São Tomé (Table S1). While maternal transmission is near perfect under standard laboratory conditions (25°C), *w*Yak transmission at relatively cool 20°C (*μ* = 0.15) is imperfect and intermediate to our low (*μ* = 0.038) and high altitude (*μ* = 0.20) field estimates. The pooled *w*Yak frequency on São Tomé in 2018 was relatively high (*p* = 0.88), suggesting most matings are between infected females and infected males. Because uninfected ova produced by infected females can be eliminated by CI in this cross (Turelli and Hoffmann 1995), we chose to pair infected females with infected males for all laboratory transmission experiments to mimic the most likely cross in nature. Therefore, it is unlikely that differences in the susceptibility of uninfected ova to CI can account for differences between our estimates of *μ* in the field and the lab. Instead, we predict that unknown factors in addition to temperature contribute to the relatively imperfect *w*Yak transmission we observe at high altitude.

Relative *w*Yak titer is reduced at 20°C compared to 25°C, suggesting temperature effects on titer contribute to variation in imperfect maternal transmission. *w*Mel titer and maternal transmission are also reduced in transinfected *Aedes aegypti* when mosquitoes are exposed to heat stress in the lab (26-40°C) and hot temperatures in the field (>39°C) (Ulrich *et al.* 2016; Ross *et al.* 2017; Foo *et al.* 2019; Ross *et al.* 2019a). Interestingly, Foo et al. (2019) found a sex-specific effect where *Ae. aegypti* exposed to heat stress produce female progeny with relatively low *w*Mel titer. This is similar to our laboratory results at 20°C and for *w*Yak variants paired with the *L42 D. yakuba* genome where transmission to female progeny was more imperfect than to males. Selection for faithful transmission to females should be relatively intense, given that the uninfected offspring produced by infected mothers may be susceptible to CI. The fact that we detected sex-biased transmission at 20°C and for the *L42* nuclear genome, but not for a yeast-enriched diet or the *L48* nuclear genome suggests biases in transmission may depend on abiotic and genetic environments.

Host diet can perturb cellular traits predicted to influence maternal transmission (Serbus *et al.* 2015; Christensen *et al.* 2019). Adult female *D. melanogaster* reared on a yeast-enriched diet for two days exhibited a 72% decrease in cellular *w*Mel titer in stage 10a oocytes (Serbus *et al.* 2015). Titer at this stage of oogenesis is predicted to determine *Wolbachia* transmission to the developing offspring (Hadfield and Axton 1999; Serbus and Sullivan 2007; Serbus *et al.* 2015; Camacho *et al.* 2017; Russell *et al.* 2018), which led us to predict that yeast-enriched food could generate imperfect transmission. Instead, maternal *w*Yak transmission is near perfect on a yeast-enriched diet (*μ* = 0.011). Despite much analysis of *Wolbachia* titer and localization in developing host oocytes (Hadfield and Axton 1999; Serbus and Sullivan 2007; Serbus *et al.* 2015; Camacho *et al.* 2017; Christensen *et al.* 2019), there have been no direct tests for covariance between these cellular traits and imperfect transmission. Our results and others (Kriesner *et al.* 2016; Ulrich *et al.* 2016; Ross *et al.* 2017; Foo *et al.* 2019; Ross *et al.* 2019a) motivate such analyses, particularly for *w*Mel-like *Wolbachia* reared at different temperatures.

### Little evidence for host or *Wolbachia* effects on imperfect transmission

*w*Yak variants in West Africa are extremely similar (0.0007% third-position pairwise differences; Cooper *et al.* 2019), having differentiated only a few thousand years ago. Indeed, *w*Yak, *w*San, and *w*Tei posess a more recent common ancestor (2500–4500 years) than do *w*Mel variants within *D. melanogaster* (4900–7200 years; Cooper *et al.* 2019). Our whole-genome analysis also suggests little differentiation between *D. yakuba* from low and high altitudes on São Tomé (mean difference = 0.0027; Figure S2). This is consistent with past genomic and phenotypic analyses of these regions by Comeault *et al.* (2016) and Turissini and Matute (2017). We found no evidence of *Wolbachia* or host effects on *w*Yak maternal transmission in our genetic analysis of reciprocally introgressed *D. yakuba*^*w*Yak^ genotypes (*L42*, *L48*, *L48*^*L42*^, *L42*^*L48*^). Interspecific crosses in *Drosophila* (Serbus and Sullivan 2007) and *Nasonia* (Funkhouser-Jones *et al.* 2018) revealed host factors that modify *Wolbachia* titer in host embryos, which could ultimately influence maternal transmission. We observed substantial heterogeneity in imperfect transmission among individual wild-caught females of *D. yakuba* and *D. santomea* ranging from *μ* = 0 to 0.929 (Figure 3), which resembles patterns observed for *w*Ri (Turelli and Hoffmann 1995; Carrington *et al.* 2011) and *w*Suz (Hamm *et al.* 2014). This could indicate that host genotypes vary in their ability to transmit *Wolbachia*, but it seems more likely that variation in host development or adult environments (e.g., temperature) underlies heterogeneity in *Wolbachia* transmission.

### The contributions of imperfect transmission, fitness effects, and CI to *Wolbachia* frequency dynamics

Our field estimates of imperfect *w*Yak and *w*San transmission do not predict *Wolbachia* frequencies on São Tomé. In particular, imperfect *w*Yak transmission was highest at high altitude *w*here *w*Yak frequency was also the highest. We estimated *μ* and *p* concurrently, assuming that *μ* does not vary significantly between host generations. If we assume that imperfect *w*Yak transmission in the *D. yakuba* generation prior to our sampling at high altitude was equal to our lowest *μ* point estimate on São Tomé (*μ* = 0.038; Table S1), we can plausibly explain the credible interval of *w*Yak frequency at high altitude with our lab estimate of weak CI (*s*_*h*_ = 0.16) and modest positive fitness effects (*F* = 1.1) (Cooper *et al.* 2017; Table S6). While it remains unknown whether *w*Yak transmission varies temporally within sites on São Tomé, three observations suggest it could. First, *w*Yak titer and *μ* are altered by temperature in the lab (Table S3). If thermal environments experienced by *D. yakuba* females in the field vary between generations, estimating *μ* in the current *D. yakuba* generation may not reflect *μ* in the prior generation. Second, we observed significant heterogeneity in *w*Yak transmission rates among *D. yakuba* females (Figure 3), suggesting *μ* may be particularly labile and depend on local conditions. Third, *w*Ri transmission varied seasonally within a single population of *D. simulans* at Ivanhoe, CA between April and November 1993 (Carrington *et al.* 2011), indicating temporal variation in *μ* within sites is possible. Future analysis of fine-scale variation in host developmental environments in the field, in concert with field and laboratory estimation of *μ*, will help elucidate the basis of *Wolbachia* frequency fluctuations.

Stronger CI and/or more positive *w*Yak effects on *D. yakuba* fitness at high relative to low altitude could potentially explain high *w*Yak frequency despite very imperfect transmission. However, even if we consider the full credible interval for *μ* at high altitude (0.087, 0.364), *F* must exceed 1.5 to approach the credible interval of *p*, regardless of CI strength (Figure S3, Table S5). Excluding the putatively anomalous YAK05b site, we still must invoke very strong CI and *F* approaching 1.5 to plausibly explain *p* using our point estimate of *μ* = 0.143. Assuming *μ* = 0.036 (the lower credible interval), our lab estimate of weak CI (*s*_*h*_ = 0.16) and *F* = 1.07 plausibly explain the lower credible interval of *p* (Figure 5, Table S5). While *w*Yak causes weak CI on average in the lab (Cooper *et al.* 2017), CI strength of some *Wolbachia* can vary across environmental conditions (Clancy and Hoffmann 1998; Ross *et al.* 2017, 2019a), host backgrounds (Reynolds and Hoffmann 2002; Cooper *et al.* 2017), and male ages (Reynolds and Hoffmann 2002). Strong positive fitness effects have not been directly estimated in this system or most others (Cooper *et al.* 2017; Shi *et al.* 2018; Meany *et al.* 2019), although *w*Ri evolved from causing a fecundity cost of *F* = 0.8–0.9 to generating a benefit of *F* = 1.1 over the course of 20 years (Weeks *et al.* 2007). Few data exist for other components of fitness, but protection from viruses and nutrient provisioning remain candidates for *w*Mel-like and other *Wolbachia* (Hedges *et al.* 2008; Teixeira *et al.* 2008; Brownlie *et al.* 2009; Osborne *et al.* 2009; Nikoh *et al.* 2014; Martinez *et al.* 2014; Newton and Rice 2020). For example, *Wolbachia* are known to provide protection from a limited number of RNA viruses under experimental conditions in the lab (Hedges *et al.* 2008; Teixeira *et al.* 2008; Osborne *et al.* 2009; Martinez *et al.* 2014), but there is currently no evidence that *Wolbachia* provide viral protection in natural *Drosophila* populations (Webster *et al.* 2015; Shi *et al.* 2018). Future work must focus on how *w*Yak and other *Wolbachia* strains benefit components of host fitness.

Given that we found no evidence that stochastic fluctuations in *w*Yak frequencies could influence the patterns we observe (Figure S4, Table S7), we predict that some combination of variable imperfect maternal transmission, *w*Yak effects on *D. yakuba* fitness, and CI strength underlie *w*Yak frequency variation on São Tomé. In contrast, spatially stable *w*San frequencies can plausibly be explained by our field estimate of imperfect transmission (*μ* = 0.068), our laboratory estimate of weak CI (*s*_*h*_ = 0.15), and modest positive fitness effects on *D. santomea*. The differences in the dynamics and equilibria of these *w*Mel-like *Wolbachia* in different host species is particularly interesting given their extreme sequence similarity across the genome.

## Conclusion

Our results add to the growing number of examples of *Wolbachia* frequency fluctuations in nature (Shoemaker *et al.* 2003; Ahrens and Shoemaker 2005; Toju and Fukatsu 2011; Hamm *et al.* 2014). Similar fluctuations have also been observed in another facultative symbiont; *Rickettsia bellii* spread to near fixation in an Arizona population of the sweet potato whitefly (*Bemisia tabaci*) in 2011, but then declined in frequency (*p* = 0.36) in 2017 (Bockoven *et al.* 2019). Our 2018 sampling on São Tomé revealed that *w*Yak frequencies vary between low and high altitudes, and *w*San varied temporally from 2015 to 2018. *Wolbachia* fluctuations are common in the *D. yakuba* clade over the last two decades (Figure 2; Cooper *et al.* 2017), and more broadly, these fluctuations seem to be a general property of *w*Mel-like *Wolbachia*. *w*Mel frequencies vary greatly among *D. melanogaster* populations across the globe (Hoffmann *et al.* 1994, 1998; Ilinsky and Zakharov 2007; Richardson *et al.* 2012; Early and Clark 2013; Webster *et al.* 2015; Kriesner *et al.* 2016). In Eastern Australia, *w*Mel frequencies decline clinally with latitude due in part to *w*Mel fitness costs in cold environments (Kriesner *et al.* 2016). Our results suggest relatively more imperfect *w*Mel transmission in cold environments could contribute to the *w*Mel frequency cline in *D. melanogaster*. Additional work evaluating the basis of spatially varying *Wolbachia* frequencies, in addition to clinal host variation (Adrion *et al.* 2015), is needed.

Understanding factors that govern *Wolbachia* spread is crucial to explain the global *Wolbachia* pandemic and to improve the efficacy of *w*Mel biocontrol (Ross *et al.* 2019b). The latter requires establishing and maintaining virus-blocking *w*Mel in mosquito-vector populations to reduce human disease transmission (e.g., dengue and Zika) (McMeniman *et al.* 2009; Hoffmann *et al.* 2011; Ross *et al.* 2019b). Our discovery that cool rearing temperature generates imperfect *w*Yak transmission opens the door to future analysis of *Wolbachia* transmission in the laboratory. Particularly promising avenues of research include the timing of infection loss during host development and the specific changes to *Wolbachia* titer and localization in host oocytes that result in imperfect maternal transmission. Ultimately, understanding how abiotic conditions and host genetic backgrounds influence all three determinants of *Wolbachia* spread—imperfect maternal transmission, fitness effects, and CI—is crucial to explain the spread and maintenance of *Wolbachia* in nature.

## ACKNOWLEDGMENTS

We thank all members of the 2018 São Tomé field crew that assisted with sampling *D. yakuba* and *D. santomea*. Tim Wheeler and Paighton Noel assisted in the lab. Dave Begun, Leonie Moyle, and two anonymous reviewers provided comments that improved our manuscript. The Cooper lab group and Aaron Comeault also provided valuable feedback. We especially thank Michael Turelli for very critical comments on an earlier draft that greatly improved our manuscript. We thank the Genomics Core and the Environmental Control for Organismal Research Laboratories at the University of Montana for their support. Research reported in this publication was supported by the National Institute of General Medical Sciences of the National Institutes of Health (NIH) under award numbers R01GM121750 to D.R.M. and R35GM124701 to B.S.C.

## SUPPLEMENTAL MATERIALS

### SUPPLEMENTAL METHODS

We selected plausible combinations of *μ*, *F*, and *H* for *w*Yak at low and high altitude based on our mathematical analyses (Table S5) to illustrate how stochasticity contributes to variation in *w*Yak frequencies at infection equilibria 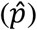. This included a set of plausible parameters for *w*Yak at low altitude (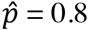, *μ* = 0.038, *s*_*h*_ = 0.16, *F* = 1.05), high altitude (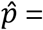 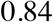, *μ* = 0.087, *s*_*h*_ = 0.20, *F* = 1.55), and a second set for high altitude with the anomalous YAK05b trap removed (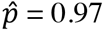, *μ* = 0.036, *s*_*h*_ = 0.45, *F* = 1.47; see Results).

Monte Carlo simulations with 10,000 replicates of population events were enacted using functions in the PopTools package (Hood 2011) and the following parameters described in Kriesner and Hoffmann (2018): (1) the number of reproductively successful females in each cohort (*N*_cs ♀_) where cohort is based on infection status, random binomial with *n* = total number of females comprising cohort (*N*_c ♀_) and *p* = 0.91, (2) the number of ova produced for each cohort, *N*_cs ♀_ × *F* × ω, where ω = normally distributed random variable with mean = 24.4 and 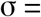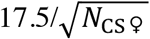, (3) the number of uninfected ova produced by infected mothers per cohort, random binomial with *n* = total number of ova produced per cohort and *p* = *μ*, and (4) the number of uninfected ova fertilized by sperm from *w*Yak-infected males and rendered inviable due to CI, random binomial with *n* = number of uninfected ova arising per cohort and *p* = *s*_*h*_. For the number of reproductively successful females and the number of ova produced for each cohort (parameters 1 and 2 above), values for *p*, mean of the normal distribution, and σ are based on fecundity data obtained previously using *D. yakuba* of varying infection status from Cooper *et al.* (2017). Here, *p* was calculated as the mean proportion of mated females in 24 hours, and the mean and σ were calculated from the daily number of eggs laid in the intraspecific *D. yakuba* fecundity experiments described in Cooper *et al.* (2017). All other parameters were set to the defaults described in Kriesner and Hoffmann (2018), including the number of viable ova per cohort that survive to adulthood, the number of successful male matings for each new cohort, and the number of reproductively successful females in each new cohort that mated with males of each infection type. All other relevant information can be found in the main text.

### SUPPLEMENTAL FIGURES

**SUPPLEMENTAL FIGURE S1.**
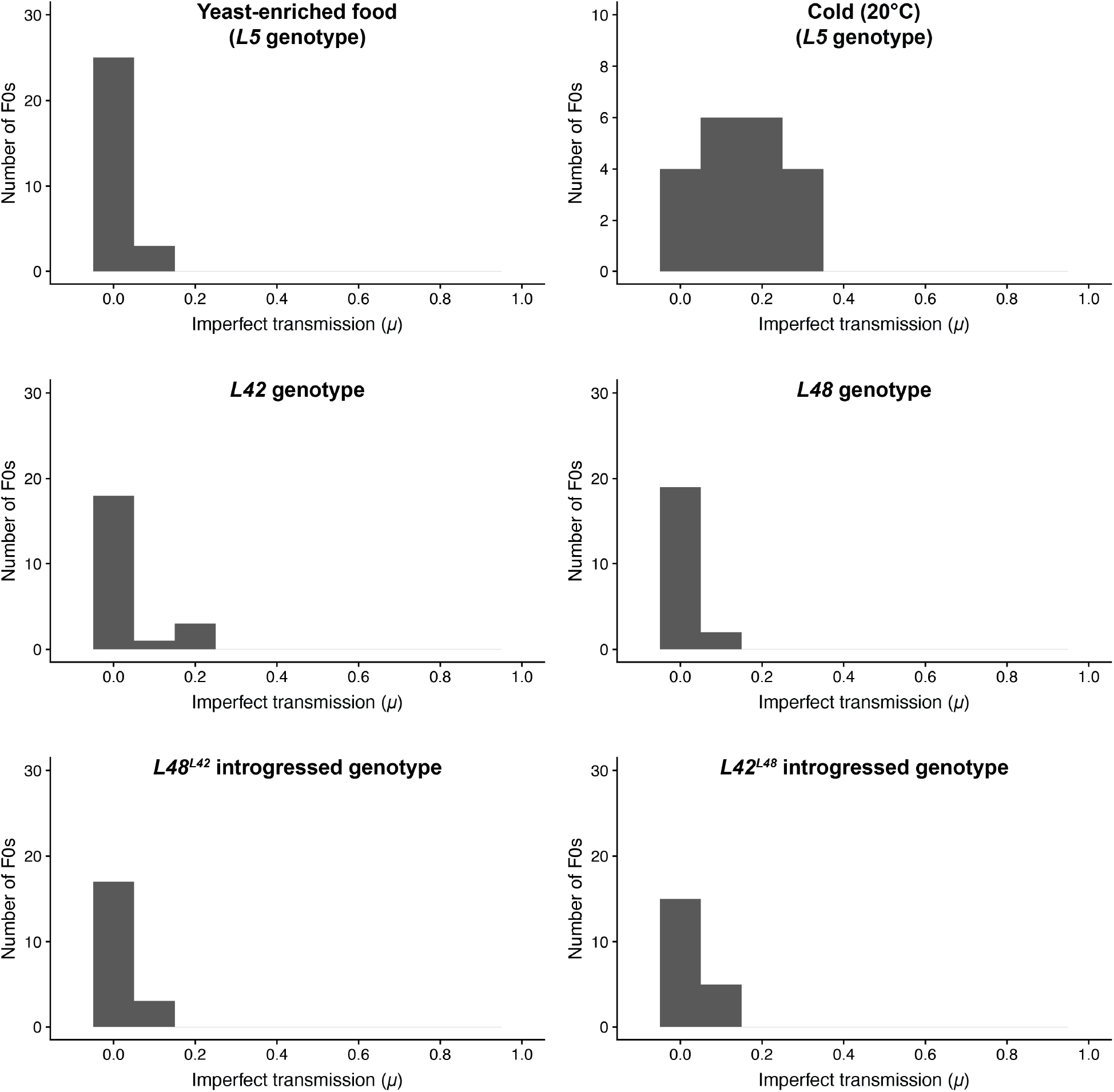
Histogram of *μ* in laboratory experiments using the *D. yakuba* isofemale line *L5* and reciprocally introgressed genotypes (*L42*, *L48*, *L48*^*L42*^, *L42*^*L48*^).

**SUPPLEMENTAL FIGURE S2.**
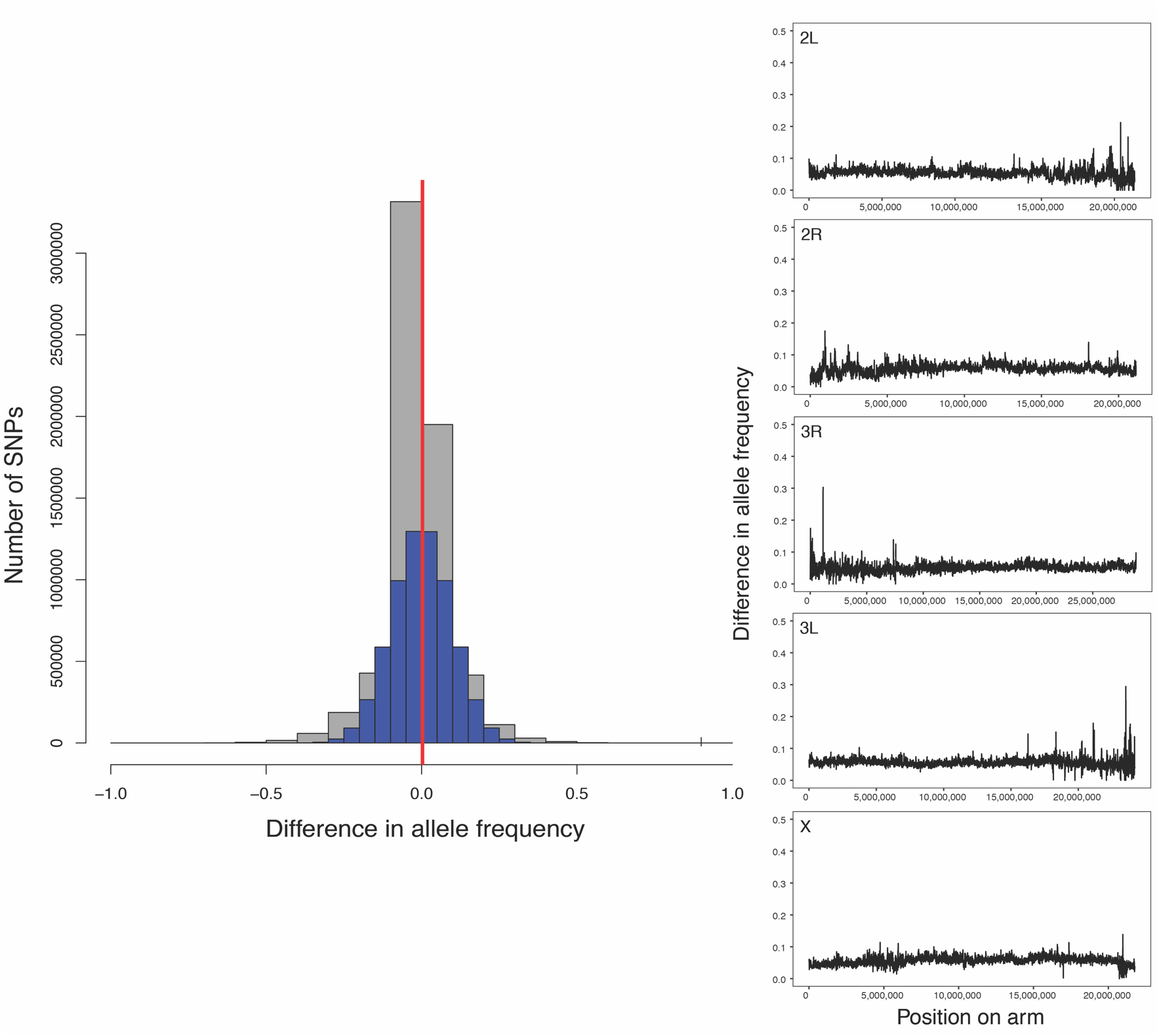
Left) Differences in allele frequencies between the genomes of *D. yakuba* from low and high altitudes on São Tomé. Histogram (gray) shows the full distribution of differences in allele frequencies between low and high altitude populations. The mean difference (0.0027) is shown with a vertical red line. This distribution overlaps the histogram of data simulated with an equivalent standard deviation and mean 0 (blue). Right) Sliding window average differences across each chromosome.

**SUPPLEMENTAL FIGURE S3.**
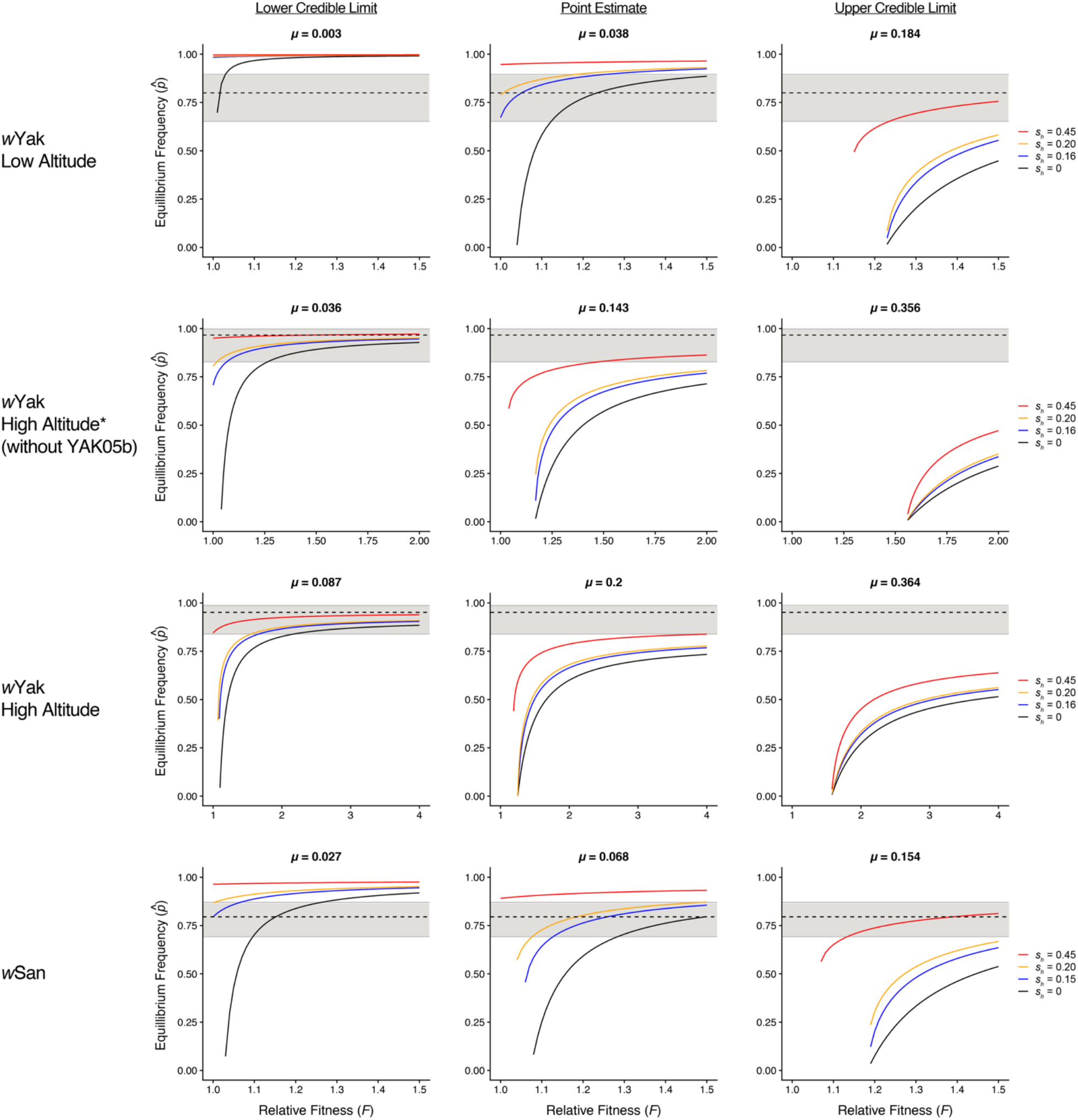
Equilibrium infection frequencies for *w*Yak and *w*San plotted against a range of *F* values, assuming our field estimates of *μ* (± credible intervals). The dotted lines indicate observed infection frequencies, and the gray areas denote 95% binomial confidence intervals. Plotted black lines denote no CI (*s*_*h*_ = 0), followed by laboratory estimates of weak CI in blue for *w*Yak (*s*_*h*_ = 0.16) and *w*San (*s*_*h*_ = 0.15) (Cooper *et al.* 2017), moderate CI in orange (*s*_*h*_ = 0.20), and strong CI in red (*s*_*h*_ = 0.45). Each row presents results for specific *Wolbachia* and regions. Plots are arranged in columns by the lower credible limit, point estimate, and upper credible limit of *μ* for each group. The asterisk signifies that the YAK05b trapping site was removed from the high altitude *w*Yak dataset due to an anomalously high value of *μ*. The full range of parameter estimates are shown in Table S5.

**SUPPLEMENTAL FIGURE S4.**
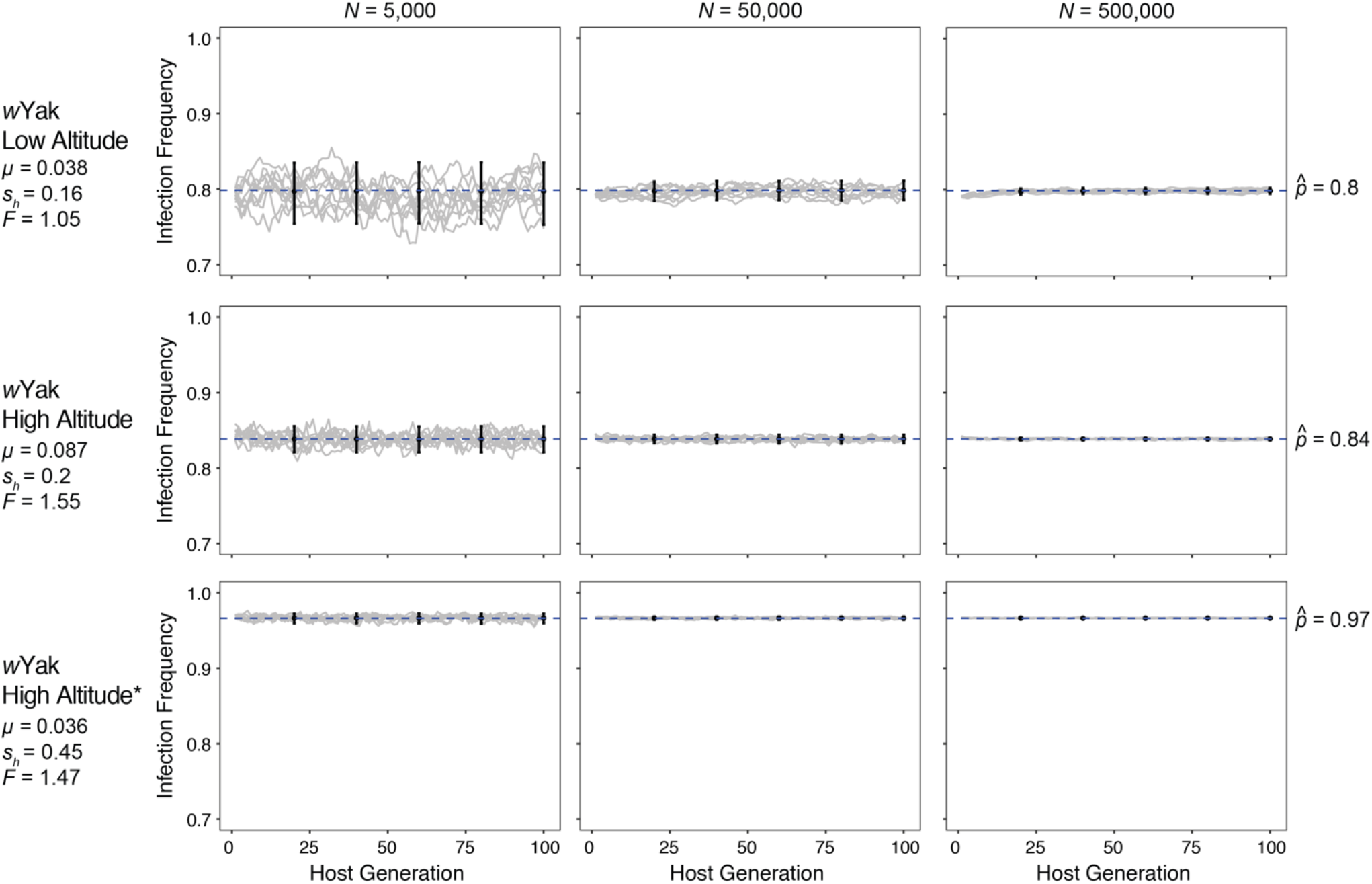
Stochastic model outcomes for *w*Yak frequencies in host populations at infection equilibrium 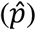. Infection dynamics over 100 host generation (10 random trials each) are shown with the mean of Monte Carlo simulations (10,000 replicates) and associated 95% confidence intervals sampled every 20 host generations. Infection equilibria predicted by Equation 2 for each region are shown with a blue dashed line. Models incorporate field estimates of *p* and *μ* and a range of *s*_*h*_ and *F* values (see Table S5). The asterisk signifies that the YAK05b trapping site was removed from the high altitude *w*Yak dataset due to an anomalously high value of *μ*. For each region, separate models were run assuming a host census population size (*N*) of 5,000, 50,000, and 500,000.

**SUPPLEMENTAL TABLE S1.** Estimates of infection frequencies (*p*) and imperfect maternal transmission (*μ*) for individual trapping sites. Sample sizes (*N*), infection frequencies (*p*) with exact 95% binomial confidence intervals, mean number of F1 offspring per family (Mean *N* F1s), and weighted mean imperfect transmission (*μ*) are shown for each trapping site. 95% BC_a_ confidence intervals were not calculated for *μ* due to inadequate sample sizes at some individual trapping sites.

| Species | Grouping | Latitude | Longitude | Elevation | Bioclim Annual<br>Mean Temperature<br>(°C) | $N$ | $N$<br>infected | $p$<br>[Confidence Interval] | Mean<br>$N$ F1s | $\mu$ |
| --- | --- | --- | --- | --- | --- | --- | --- | --- | --- | --- |
| *D. yakuba* | Low Altitude |  |  |  |  |  |  |  |  |  |
|  | YAK02 | 0.304 | 6.646 | 590 | 20.9 | 40 | 32 | 0.800 [0.652, 0.895] | 9.9 | 0.038 |
|  | High Altitude |  |  |  |  |  |  |  |  |  |
|  | YAK04 | 0.290 | 6.630 | 900 | 20.6 | 27 | 26 | 0.963 [0.817, 0.998] | 10.6 | 0.148 |
|  | YAK05 | 0.289 | 6.616 | 1096 | 17.8 | 2 | 2 | 1.000 [0.342, 1.000] | 8.5 | 0.059 |
|  | YAK05b | 0.290 | 6.616 | 1104 | 17.8 | 12 | 11 | 0.917 [0.646, 0.996] | 9.7 | 0.354 |
| *D. santomea* | Site 1 |  |  |  |  |  |  |  |  |  |
|  | CAR01 | 0.277 | 6.588 | 1184 | 17.8 | 3 | 2 | 0.667 [0.208, 0.983] | 14.3 | 0.133 |
|  | CAR02 | 0.277 | 6.588 | 1216 | 17.8 | 3 | 3 | 1.000 [0.439, 1.000] | 5.0 | 0.000 |
|  | CAR03 | 0.277 | 6.586 | 1261 | 17.8 | 3 | 2 | 0.667 [0.208, 0.983] | 6.0 | 0.308 |
|  | CAR04 | 0.279 | 6.587 | 1347 | 17.8 | 4 | 2 | 0.500 [0.150, 0.850] | 8.0 | 0.000 |
|  | CAR05 | 0.279 | 6.589 | 1340 | 17.8 | 11 | 10 | 0.909 [0.623, 0.995] | 11.9 | 0.130 |
|  | Site 2 |  |  |  |  |  |  |  |  |  |
|  | CAR07 | 0.279 | 6.594 | 1414 | 17.8 | 9 | 8 | 0.889 [0.565, 0.994] | 9.1 | 0.000 |
|  | CAR09 | 0.281 | 6.595 | 1399 | 17.8 | 8 | 5 | 0.625 [0.306, 0.863] | 8.3 | 0.000 |
|  | CAR10 | 0.282 | 6.596 | 1411 | 17.8 | 16 | 12 | 0.750 [0.505, 0.898] | 8.6 | 0.000 |
|  | TRNE01 | 0.281 | 6.592 | 1460 | 17.8 | 10 | 8 | 0.800 [0.490, 0.943] | 12.5 | 0.171 |
|  | TRNE02 | 0.282 | 6.593 | 1464 | 17.8 | 7 | 7 | 1.000 [0.646, 1.000] | 11.4 | 0.000 |
|  | TRNE03 | 0.283 | 6.597 | 1385 | 17.8 | 4 | 3 | 0.750 [0.301, 0.987] | 8.8 | 0.033 |

**SUPPLEMENTAL TABLE S2.**
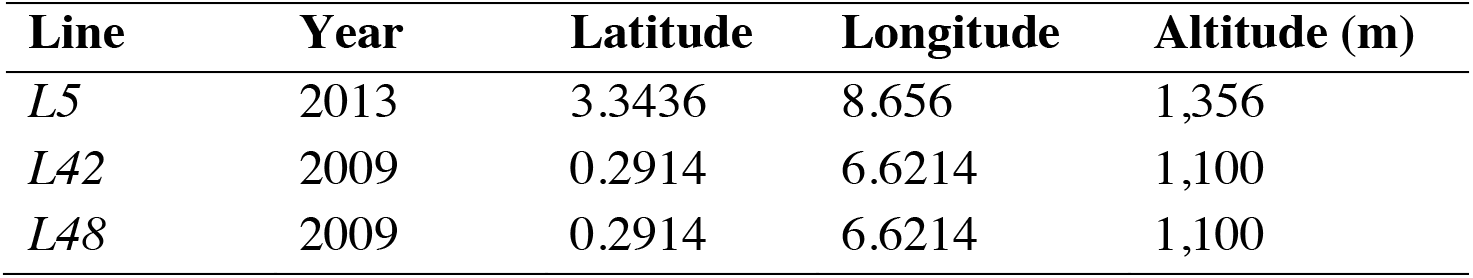
Sampling information for isofemale lines used in laboratory experiments.

**SUPPLEMENTAL TABLE S3.** Imperfect *w*Yak maternal transmission in the laboratory. The number of sublines (*N*), mean number of F1 offspring per subline (Mean *N* F1s), weighted mean imperfect transmission (*μ*), and 95% BC_a_ confidence intervals are shown for each experiment.

| **Treatment** | **$N$** | **Mean $N$ F1s** | **$\mu$ [Confidence Interval]** |
| --- | --- | --- | --- |
| yeast-enriched food | 28 | 9.89 | 0.011 [0.004, 0.029] |
| cold (20°C) | 20 | 10 | 0.15 [0.109, 0.196] |
| host genotype *L42* | 22 | 10 | 0.032 [0.009, 0.082] |
| host genotype *L48* | 21 | 9.86 | 0.01 [0, 0.034] |
| reciprocal introgression *L42*<sup>*L48*</sup> | 20 | 9.95 | 0.025 [0.01, 0.055] |
| reciprocal introgression *L48*<sup>*L42*</sup> | 20 | 10 | 0.015 [0.005, 0.045] |

**SUPPLEMENTAL TABLE S4.** Gene annotations for completely differentiated SNPs between low and high altitude populations of *D. yakuba*. Annotation results are shown each SNP using the *D. yakuba* Release 2.0 assembly and annotation on the UCSC Genome Browser.

| Chromosome Arm | SNP Position | Position | Annotation Gene Symbol | Dmel Gene Symbol | Gene Name | Gene Location | Molecular Function (GO) Predictions |
| --- | --- | --- | --- | --- | --- | --- | --- |
| 3L | 3668737 | <10kb upstream | CG7213 | CG7213 | NA | chr3L:3,668,973-3,672,288 | NA |
|  |  | <10kb upstream | CG8023 | eIF4E3 | eukaryotic translation initiation factor 4E3 | chr3L:3,673,908-3,674,928 | RNA 7-methylguanosine cap binding; RNA cap binding; contributes to translation initiation factor activity; translation initiation factor binding |
| 3L | 5201729 | <10kb upstream | CG7471 | HDAC1 | Histone deacetylase 1 | chr3L:5,202,077-5,204,254 | histone deacetylase activity; transcription corepressor activity |
|  |  | <10kb upstream | CG43225 | axo | axotactin | chr3L:5,205,034-5,205,411 | serine-type endopeptidase inhibitor activity |
|  |  | <10kb upstream | CG13716 | NA | NA | chr3L:5,209,817-5,210,164 | NA |
| 3L | 18327412 | 10kb intergenic | NA | NA | NA | NA | NA |
| 3R | 16902666 | <10kb upstream | CG18765 | NA | NA | chr3R:16,904,248-16,905,597 | NA |
|  |  | <10kb upstream | CG6908 | NA | NA | chr3R:16,906,043-16,907,540 | NA |
|  |  | <10kb upstream | CG6834 | NA | NA | chr3R:16,907,939-16,911,105 | NA |
|  |  | <10kb upstream | CG6830 | NA | NA | chr3R:16,911,662-16,914,246 | NA |
| 3R | 16902675 | <10kb upstream | CG18765 | NA | NA | chr3R:16,904,248-16,905,597 | NA |
|  |  | <10kb upstream | CG6908 | NA | NA | chr3R:16,906,043-16,907,540 | NA |
|  |  | <10kb upstream | CG6834 | NA | NA | chr3R:16,907,939-16,911,105 | NA |
|  |  | <10kb upstream | CG6830 | NA | NA | chr3R:16,911,662-16,914,246 | NA |
| X | 3902534 | <10kb upstream | CG3062 | NA | NA | chrX:3,905,462-3,906,454 | NA |
|  |  | <10kb upstream | CG3081 | NA | NA | chrX:3,906,928-3,908,664 | NA |
|  |  | <10kb upstream | CG3346 | pon | partner of numb | chrX:3,909,330-3,911,475 | NA |
|  |  | <10kb upstream | CG12179 | NA | NA | chrX:3,912,017-3,915,981 | NA |
| X | 5614370 | 10kb intergenic | NA | NA | NA | NA | NA |
| X | 8177508 | <10kb upstream | CG18319 | ban | bendless | chrX:8,186,055-8,186,507 | ubiquitin conjugating enzyme activity |

**SUPPLEMENTAL TABLE S5.**
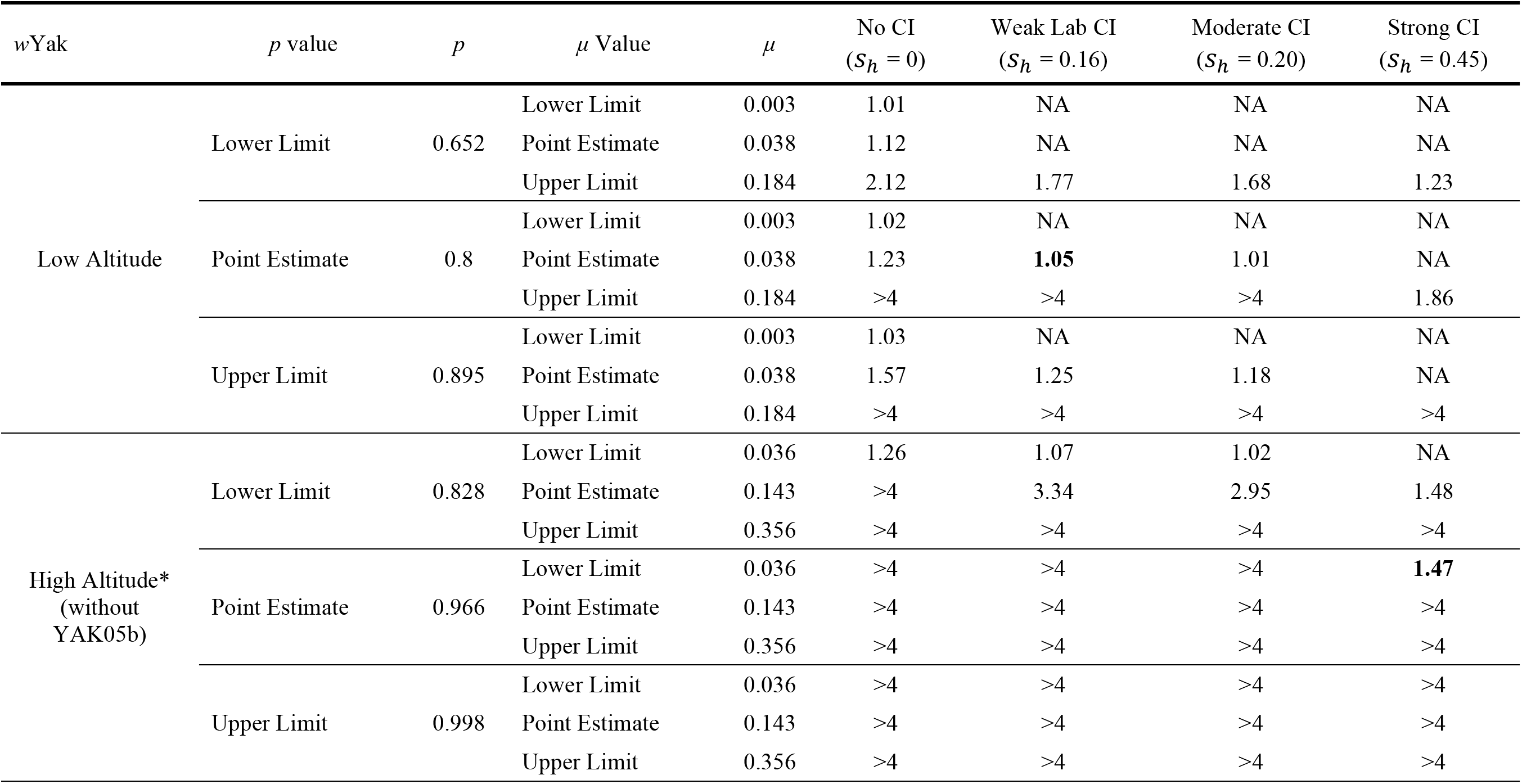
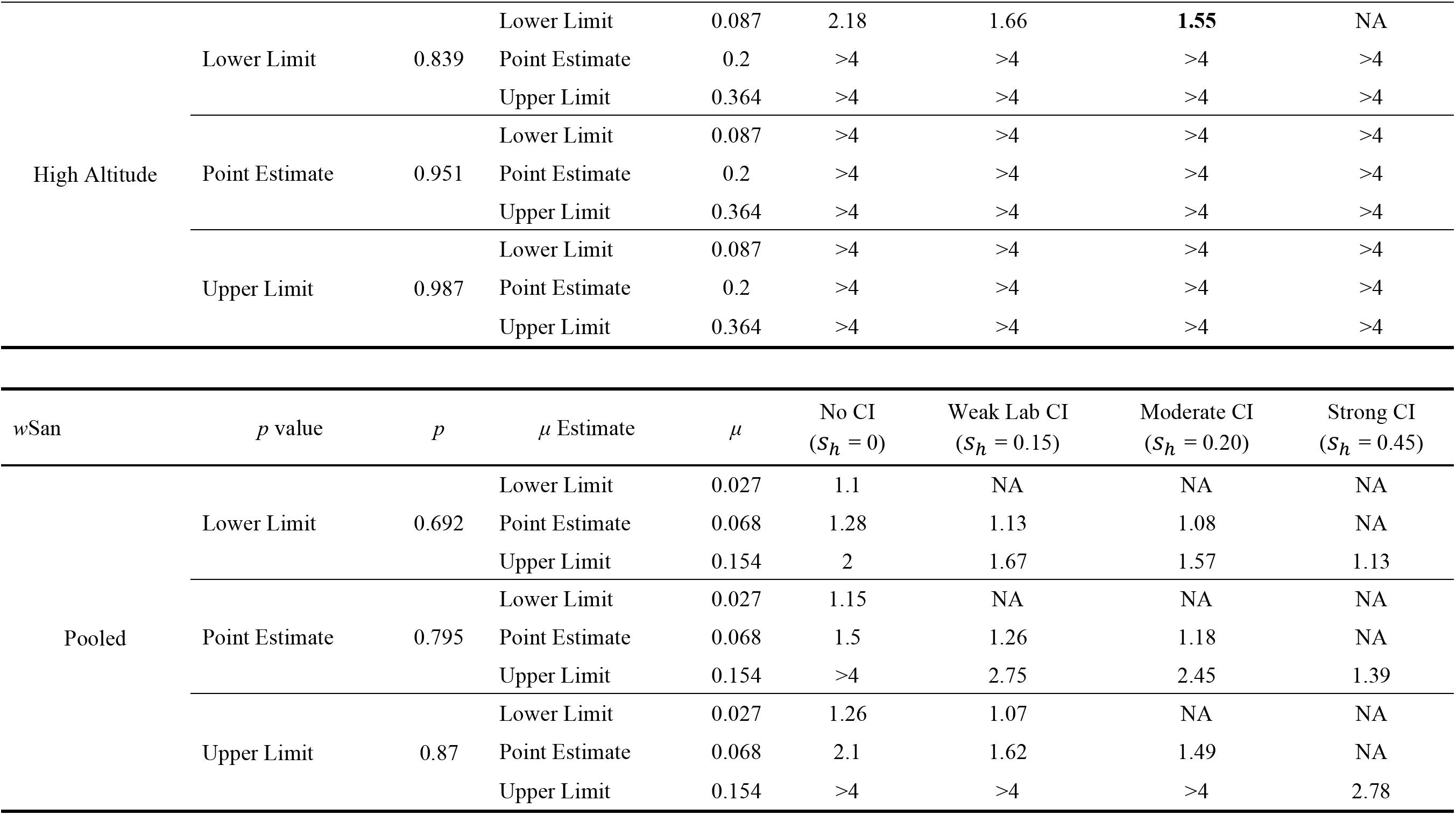
Values of *F* required to explain estimates of *p* (± credible intervals) given our field estimates of *μ* (± credible intervals) and a range of CI strength. Estimates were calculated using equation 2. Estimates of infection frequencies (*p*) and weighted mean imperfect transmission (*μ*) are shown for *w*Yak at low and high altitude sites and for the pooled sample of *w*San. For each grouping, we evaluated parameter space across the full credible interval of *p* and *μ* (lower limit, point estimate, and upper limit). Cells labeled NA indicate bistable equilibria where *F*(1 – *μ*) < 1, which we consider unlikely because they preclude *Wolbachia* spread at low frequency (counter to observations in nature; see Methods). Values of *F* > 1.5 are considered biologically unrealistic and have not been observed in any system (see Meany *et al.* 2019). *F* values in bold identify parameter combinations used the stochastic models adapted from Kriesner and Hoffmann (2018) (Figure S4, Table S7).

**SUPPLEMENTAL TABLE S6.**
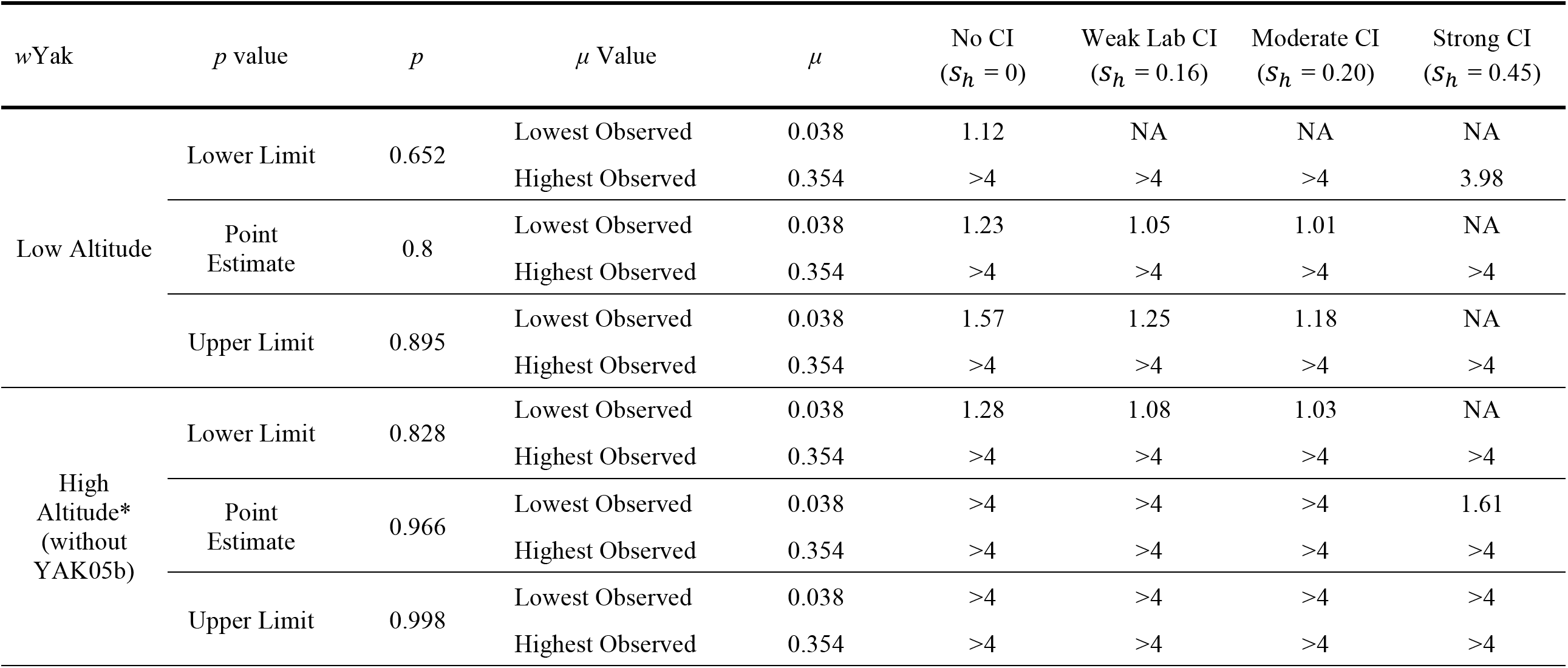
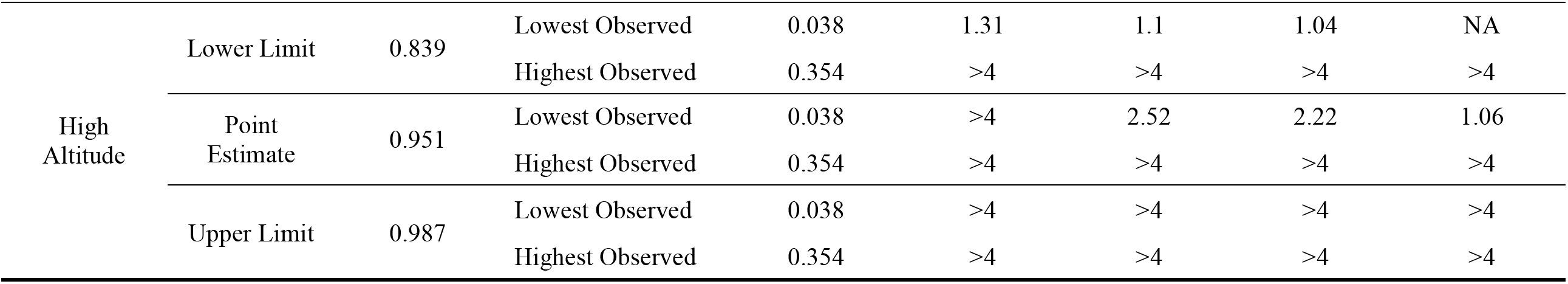
Exploration of parameter space assuming *w*Yak *μ* in the current generation may not reflect *μ* in the previous generation. We conservatively considered the full range of *w*Yak *μ* point estimates (0.038–0.354) across all *D. yakuba* trapping sites on São Tomé in our 2018 sample (Table S1), which exceeds seasonal variation of *w*Ri *μ* observed in a population of *D. simulans* in Ivanhoe, CA (Turelli and Hoffmann 1995). We assume that any between-host-generation variation in *μ* within a region is unlikely to exceed the full range of *μ* point estimates across all trapping sites (Table S1). Values of *F* required to explain estimates of *p* (± credible intervals) for each group were calculated using equation 2. Cells labeled NA indicate bistable equilibria where *F*(1 – *μ*) < 1, which we consider unlikely because they preclude *Wolbachia* spread at low frequency (counter to observations in nature; see Methods). Values of *F* > 1.5 are considered biologically unrealistic and have not been observed in any system (see Meany *et al.* 2019).

**SUPPLEMENTAL TABLE S7.**
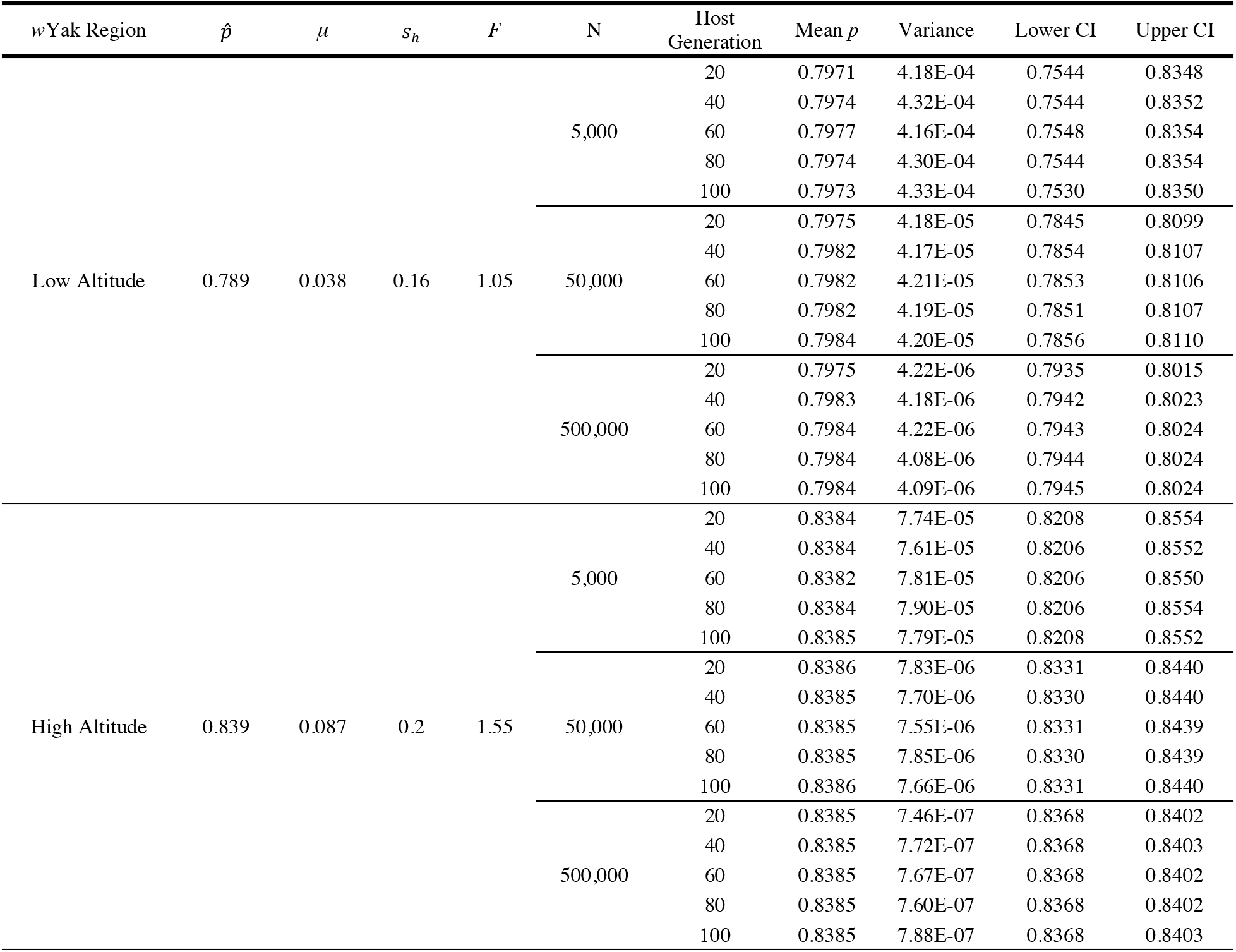
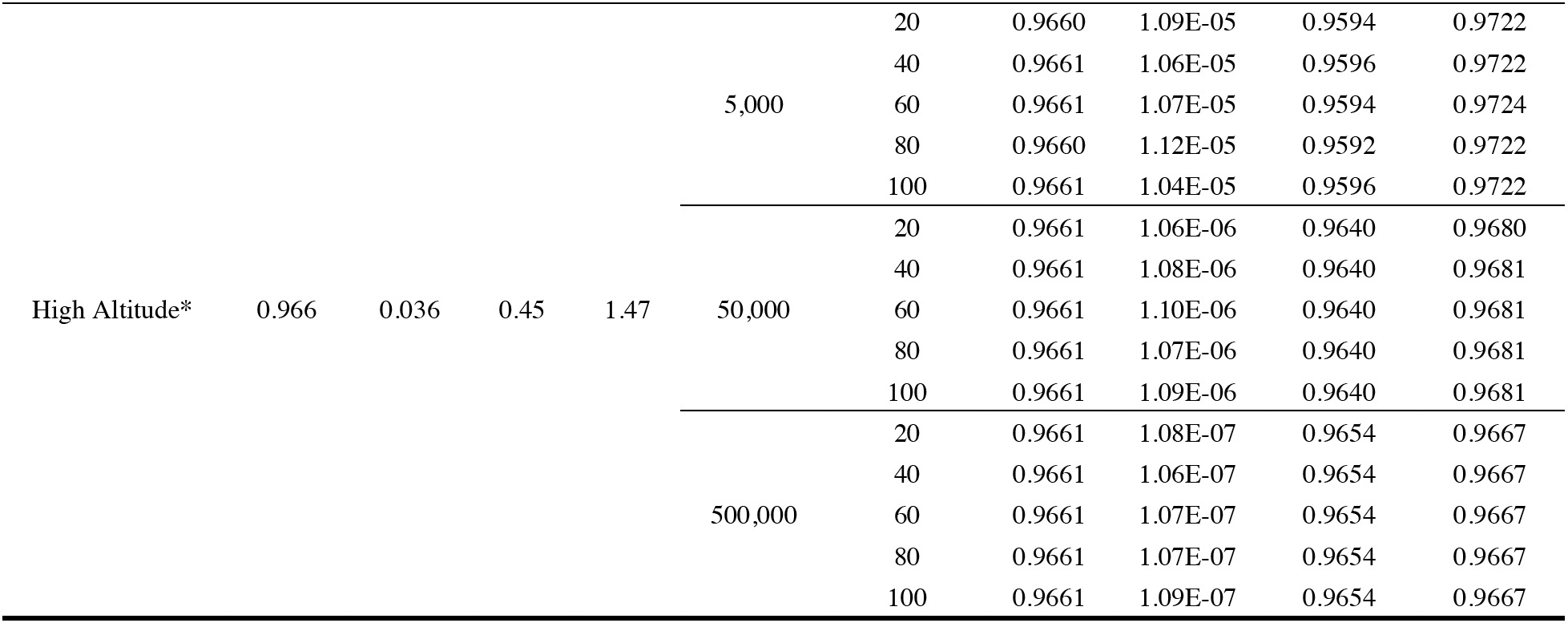
Stochastic model outcomes for *w*Yak frequencies in populations at infection equilibrium 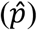. Models assume field estimates of 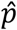 and *μ* and plausible values of *s*_*h*_ and *F* for each region (see Table S5). The asterisk signifies that the YAK05b trapping site was removed from the high altitude *w*Yak dataset due to an anomalously high value of *μ*. For each region, separate models were run assuming a host census population size (*N*) of 5,000, 50,000, and 500,000. Infection dynamics over 100 host generation are shown with the mean infection frequencies (*p*), variance, and associated 95% confidence intervals from the Monte Carlo simulations (10,000 replicates).

**SUPPLEMENTAL TABLE S8.**
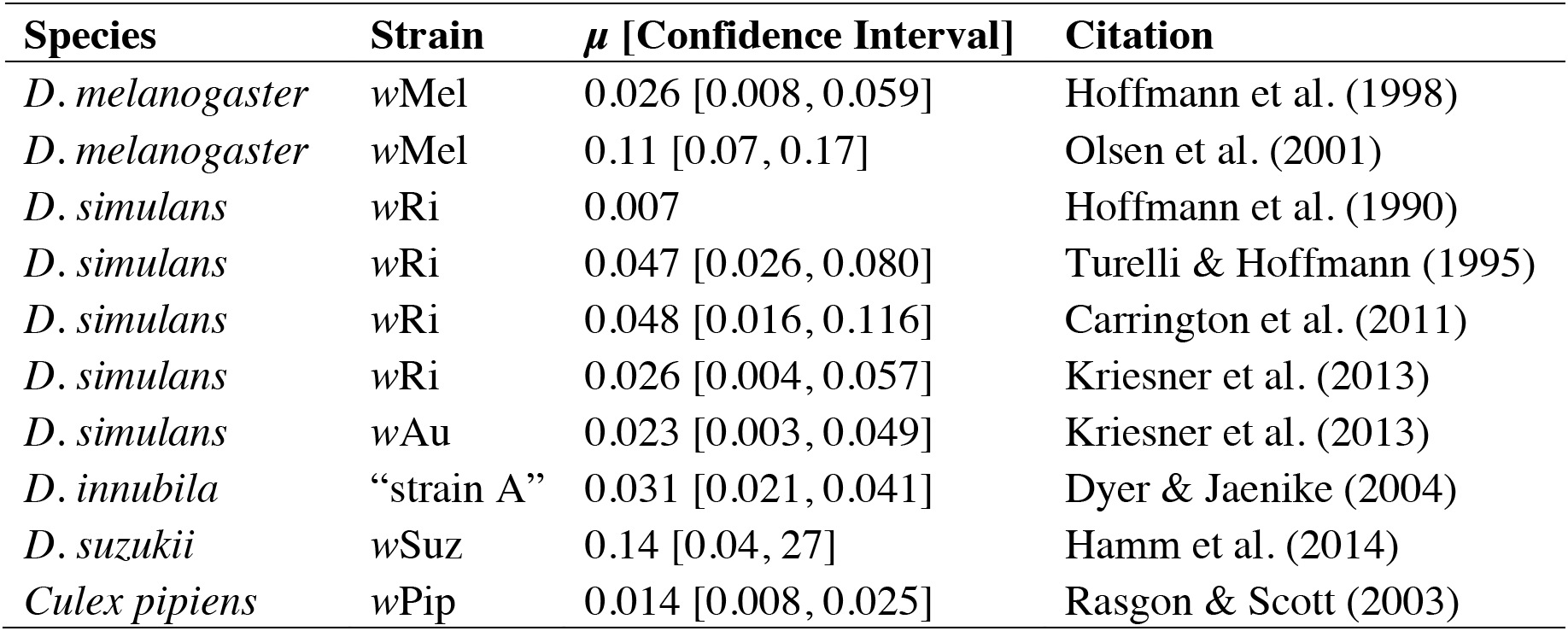
Estimates of imperfect maternal transmission from wild-caught females. Estimates of *μ* and 95% confidence intervals (when provided) are shown from previous work. See citations for specific details on *μ* estimation for each study.

